# Graph attention with structural features improves the generalizability of identifying functional sequences at a protein interface

**DOI:** 10.1101/2025.11.04.686550

**Authors:** J. Ash, I. M. Francino-Urdaniz, S. P. Kells, C. N. Davis, T. A. Whitehead, S. D. Khare

## Abstract

Accurate prediction of the set of sequences compatible with a protein-protein interface is an unsolved problem in biology. While supervised sequence-based models trained directly on experimental data can predict variant effects, they often fail to generalize to significantly diverged sequences. We hypothesized that incorporating information from deep learning models of proteins (e.g., ESM, ProteinMPNN) could enhance generalizability. To test this hypothesis, we designed and experimentally screened several deep mutational libraries of the SARS-CoV-2 Spike Receptor Binding Domain (RBD) for binding to the ACE2 receptor. Our large dataset encompasses over 43,000 sequence variants, exhibiting up to 26 substitutions away from the parental RBD sequence, thus exploring a significantly expanded sequence space compared to previous studies. Baseline supervised learning with one-hot encoded sequences achieved high accuracy within training sets but poor performance on unseen libraries. Integrating pre-trained protein model embeddings (ESM2) as a feature showed modest improvement in generalization. To further enhance predictive power, we developed a graph attention network architecture that combines representations of local residue environments using protein structure graphs with long-range inter-residue correlations captured by protein language model (PLM) embeddings (GAN-PLM). By explicitly modeling residue environments, interface geometry, and sequence dependencies, our graph attention model outperformed purely sequence-based models, achieving substantially higher balanced accuracies when predicting functional ACE2-binding variants across the diverse sequence space spanned by our independent libraries. This demonstrates the potential of structure- and sequence-based features into deep learning frameworks to achieve accurate and generalizable predictions of protein interface function, with broad implications for understanding and engineering protein interactions relevant to emerging infectious diseases and therapeutic protein design.

## Introduction

Evolution shapes the sequence variation at protein-protein interfaces. On short evolutionary timescales, *e.g.*, for pathogen interactions with proteins exploited for cell entry, sequence variation occurs on only one side of the interface. An open question in biology is the allowable sequence variation of one side of a protein interface. Since a classical analysis from Wells ^1^, researchers have found that relatively limited experimental data can predict the function of a large fraction of observed functional variant effects using interpretable supervised models. These models largely consider residue preferences, with only a minor epistatic contribution between residues ^2,3^. On the one hand, such models are powerful in that high throughput experimental binding data for new protein interactions can quickly be assessed to guide prediction. On the other hand, such models cannot generalize beyond the narrow confines of the variation at specific residue positions in a particular protein. These models will fail when the sequence of the interface changes considerably, and cannot be used for accurate zero shot predictions. Such predictions could be useful for prognosticating future outbreaks of infectious diseases, for understanding the functional basis of affinity and specificity within naturally occurring and *de novo* designed binding proteins, or for designing plasticity and robustness in protein-ligand recognition more broadly.

Physics-based models are general but imprecise, with a mean unsigned error in the range of 0.5-1 kcal/mol per mutation. Given that a 10-fold reduction in binding affinity is 1.4 kcal/mol at 37°C, physics-based models quickly lose predictive power extending beyond a handful of mutations at a given interface. Recent advances in deep learning for biophysical problems like structure prediction from sequence ^4,5^ suggests that deep learning approaches have the potential to predict this set of allowable sequences in a generalizable manner. Foundational models like ProteinMPNN ^6^ and ESM ^5^ have learned what sequences encode a viable protein. In principle, favorable sequences inferred from these models could lead to identifying the complete set of functional sequences at a protein interface.

We hypothesized that, with the recent deep learning advances in protein design, we could design, measure, and predict very large shifts in protein sequence space by enriching mutational libraries with functional sequences. To test this prediction, we designed several deep mutational libraries of the SARS-CoV-2 Spike receptor binding domain (RBD), and assessed library members for maintenance of ACE2 binding using yeast surface display coupled to fluorescence activated cell sorting. Functional binding variants contain up to 23 mutations away from the parental Omicron strain and show considerable plasticity of RBD sequence variation. Consistent with previous studies, we found that one hot encoding of sequence features can forecast the ACE2 binding of individual sequences, but that these predictions do not extend to sequences outside of the training set. Geometry-aware graph-based approaches combined with protein language models ^5^ captures both representations of local residue environments and interface geometry via graph representations and long-range sequence dependencies via protein language models (ESM-2), resulting in significant increases in generalized classification accuracy compared with purely sequence-based models. The present combination of high-throughput binding screening with deep sequencing and geometric deep learning has the potential to allow unprecedented insight into plasticity at protein-protein interfaces.

## Results

### Deep mutagenesis and screening of RBD ACE2-binding sequences

The SARS-CoV-2 S RBD:ACE2 interaction provides a good testbed for developing and testing the ability of various methods to predict allowable sequence variation. Earlier efforts at assessing RBD sequence diversity focused largely on the local mutational landscape (1-2 mutations away from the parental sequence). Starr et al. developed a yeast display platform for evaluating the sequence-function landscape of Wuhan Hu-1 S RBD by deep mutational scanning ^7^. This landmark paper was followed by efforts from multiple groups at evaluating sequence tolerance for ACE2 recognition and/or propensity for escape from antibody-mediated virus neutralization ^8–10^. Reddy et al. ^11^ sampled RBD at more depth but still within the confines of natural selection observed by Omicron and later VOCs ^12,13^. More recently, Bahl and colleagues computationally identified functional RBD sequences distal in sequence space, and tested a few dozen sequences^14^. To sample the functional binding space more broadly, we constructed several sequence-binding experimental datasets that represent large shifts in sequence space from an Omicron chimeric S RBD (S RBD (333-541) Wuhan-1 with mutations K417N, S477N, E484A, Q498R, N501Y and Y505H). We denoted this Omicron chimeric S RBD wild-type or ‘WT’ as it was the starting point for all libraries. In total, we designed and assembled four separate RBD mutational libraries covering the solvent-accessible positions of the Barnes and Bjorkman-defined Class 1 or Class 2 epitopes around the receptor binding site (RBS) ^15^. Our combined libraries contain 43,330 variants with multiple mutations at 42 distinct positions in the Class I and/or Class II epitopes. Libraries were assembled into a yeast display plasmid and assessed by yeast surface display coupled to Fluorescence Activated Cell Sorting (FACS; **Fig 1A**). ACE2-binding properties of yeast displayed S RBD variants correlate well with ACE2 binding of full-length S *in vitro*, as demonstrated by multiple groups ^7,8–10,11^.

**Figure 1.**
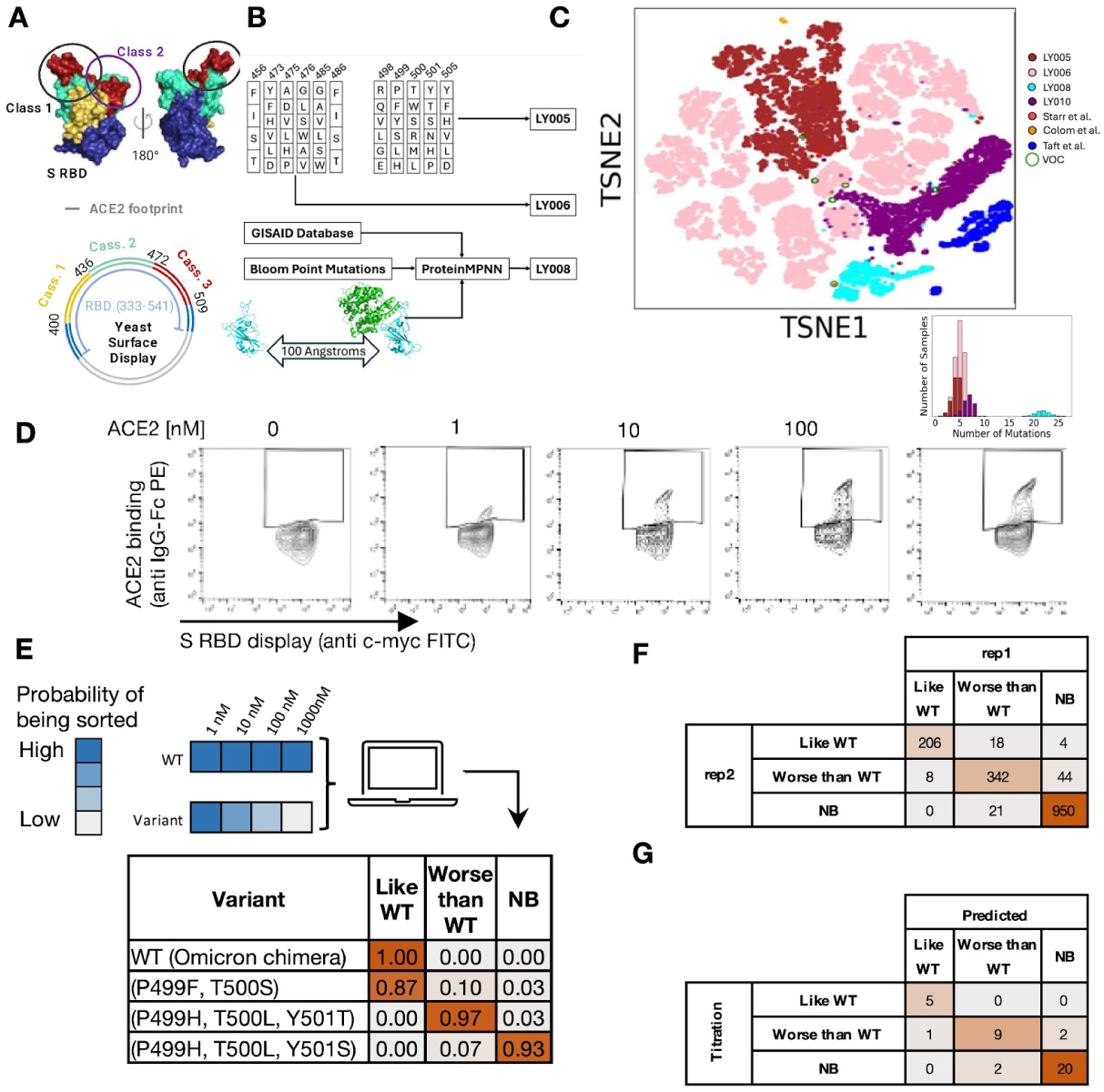
Design, construction, and testing of diverse S RBD variants using yeast surface display. **A.** S RBD structure showing the division of the ACE2 footprint in 3 sequential ‘cassettes’ by color code. The Barnes et al. ^15^ epitope classification is circled on the structure. The yeast display plasmid map indicating cassette boundaries (cass. 1-3) is displayed at the bottom of the panel. The colors correspond to the encoded structural positions in the structure shown in panel A. Portions of the protein that are unmutated are shown in blue on the structure and plasmid. **B.** Schematic of the design process for the initial three libraries LY005, LY006, LY008. **C.** 2D t-SNE^20^ plot showing VOCs, previous experimental datasets^7,11,14^, and libraries generated in this work. **D.** Collection gates for binding variants on ACE2 binding vs RBD display cytograms at 0nM to set the gate above noise, as well as collecting gates at 1nM, 10nM, 100nM and 1000nM ACE2. **E.** The probability of sorting at each labeling concentration is converted to one of three classifications: binding to S RBD “like WT”, “worse than WT” or “no binding”. This classification takes the form of a probability vector. The inset table shows classification of representative variants. **F.** Replicate sorts of library LY008 (rep1, rep2) showing the consistency of the experiment. **F.** Consistency of binding classification between titrations of individual variants (Titration) and predicted classification by deep sequencing (Predicted).

The regions of the RBD targeted for design were constrained between positions 436-472 (**Fig 1A**) and 473-509. Two libraries (LY005, LY006) were encoded by degenerate codons ^16,17^ chosen to maximize sequence diversity while preserving the physicochemical basis of the natural amino acids (charged, neutral, large, small, hydrophobic, polar) (**Fig 1B**). The library LY008 was designed computationally using distinct approaches. A subset of the library was mutated at positions making direct interactions with ACE2 (positions 449, 453, 455, 456) using ProteinMPNN ^6^. The remaining library members were also designed by ProteinMPNN ^6^ biased by the Bloom dataset identifying affinity-enhancing mutations ^7^ and/or the GISAID database ^18^ identifying individual mutations observed in the population (**Methods**). Combined, these initial libraries contained a median of 6 mutations (range 1-26) (**Fig 1C**) and largely map a different sequence space than has previously been assessed naturally or synthetically (**Fig 1C**).

Our custom libraries were constructed from short DNA fragments by Golden Gate assembly ^19^ and transformed into yeast. After induction of S RBD surface expression, libraries were sorted at various ACE2 concentrations (range 1-1000 nM) using FACS. At each concentration, we collected all cells maintaining binding to ACE2 (**Fig 1D, Fig S1**) and deep sequenced the population along with a reference population. This experimental pipeline enabled a qualitative analysis of ACE2 binding, compared to WT, by correlating the probability of being sorted at each ACE2 concentration to three classification bins: “like WT”, “worse than WT” and “no binding” (**Fig 1E**; see **METHODS**).

To test internal reproducibility, we retransformed the LY008 library into yeast and sorted it on a different day (**Fig 1F**). We observed 93% agreement between biological replicates. Notably, only 0.25% of variants were classified as ‘like WT’ in one replicate and ‘no binding’ in the other replicate. To test whether titrations of individual clones correlated with population measurements, we selected 39 variants across all libraries and performed clonal titrations in replicate (**Figure S2-3**). We classified variants binding worse than WT if they had at least a 10-fold decrease in K_D_ relative to WT and at least a 2-fold increase in fluorescence over background at 1 micromolar ACE2. We observed 87.2% (34/39) agreement between clonal titrations and population measurements (**Fig 1F, Table S1**), showing reasonable agreement between measurements. To test for the possibility that RBD designs bind ACE2 non-specifically, we assessed the ability of lysozyme to bind the RBD library. While lysozyme binds non-specifically to cells, the fluorescence of the displaying population does not increase relative to the non-displaying cells indicating that it does not bind to the displayed RBD (**Fig S4**). Overall, our libraries can assess ACE2 binding of large libraries with reasonable accuracy and internal reproducibility.

### Baseline supervised learning informs the generalizability of interface plasticity prediction models

Previous work has shown that in some cases, a large fraction of the observed combinatorial variant effects on protein-protein binding can be captured by simple, interpretable supervised models that solely consider site-wise amino acid preferences (via one-hot encoding) with only a minor contribution from interactions between residues ^21^. In other studies, epistatic mutational effects were found to be the key drivers of protein function, such that the combinatorial effects of mutations were highly context dependent^22,23^. However, these models are system- and residue-position-specific, and cannot be generalized. We gauged the performance of a variety of supervised machine learning algorithms using a one-hot encoded representation of experimentally evaluated RBD sequences. We chose Support Vector Classifier, RandomForest, K-Nearest Neighbors, Naive Bayes, and Logistic Regression as our baseline architectures ^11^. We examined libraries LY005, LY006, and LY008 separately, and tasked each model to predict the activity of the holdout set across five stratified folds (**Fig. 2A**). We found that logistic regression consistently performs the best across all three datasets, achieving validation balanced accuracies (**Fig. 2B**) of 88.4%, 63.1%, and 91.9% on libraries LY005, LY006, and LY008, and Matthews Correlation Coefficient (MCC; **Fig. 2C**)^24^ values of 0.5, 0.49, and 0.74 respectively. A full set of statistical performance metrics including AUC, AUPR, and F1-score is reported in **Table S2**.

**Figure 2.**
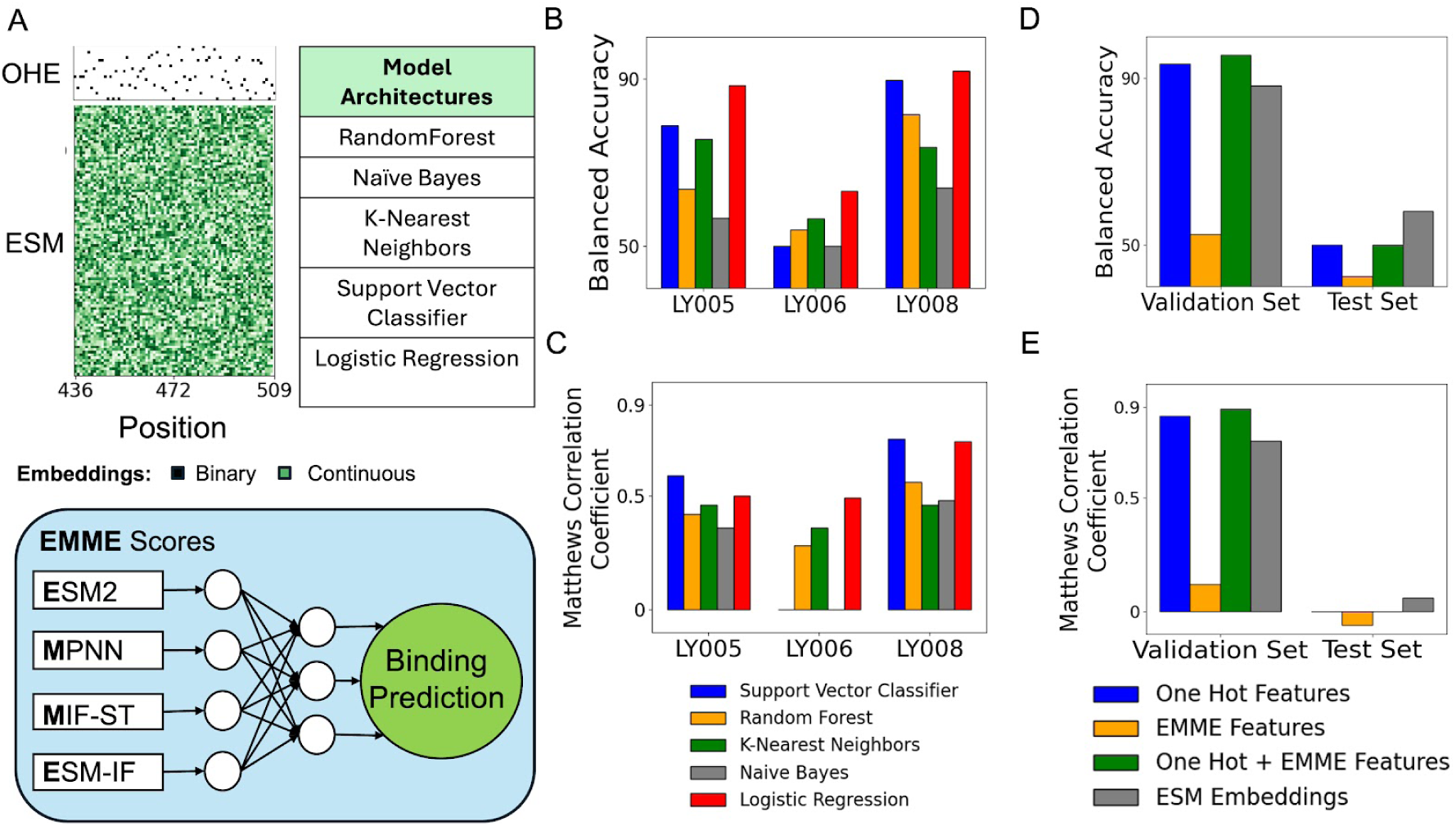
Supervised learning accurately predicts performance within but not between experimental datasets. **A**: Model architectures and features used for benchmarking RBD binding recapitulation. Graphical representations for the ESM and the one-hot encoding (OHE) embedding matrices that served as model inputs are shown along the top of the panel. The components making up the EMME scores, as well as a list of all simple architectures used, are also included at the bottom of the panel. **B, C**: Bar graphs representing the best stratified 5-fold cross validation balanced accuracies (B) and Matthews’ Correlation Coefficients (C) attained by models on each sequence library using only one-hot encoded representations. **D, E**: Prediction performance for stratified 5-fold cross validation and test set balanced accuracy (D) and Matthews’ Correlation Coefficients (E) attained using Logistic Regression with various feature representations and combinations.

Having established Logistic Regression as the best-performing architecture for recapitulating binding based on one-hot encoded sequences as features, we asked if a more transferable set of features could improve generalizability using this framework. We therefore tested a selection of score-based features derived from the pre-trained protein models <u>E</u>SM2 ^5^, Protein<u>M</u>PNN ^6^, <u>M</u>IF-ST ^25^, and <u>E</u>SM-IF ^26^ (referred to as EMME features). We also used the ESM2-derived sequence embeddings directly as a separate set of features. To investigate generalizability, we trained a logistic regression model using different combinations of EMME, one-hot encoded, and ESM2 embedding inputs for 80% of the sequences in LY008 across five stratified splits. We use this model to evaluate prediction accuracy for both the holdout set (LY008) and all of the LY005 and LY006 libraries. While EMME features were not predictive of validation or test set binding labels by themselves, they did modestly improve recapitulation when provided to the model alongside the one-hot encoded sequences. The model achieved 93.5% balanced validation accuracy using only one-hot encoded predictors, which increased to 95.6% with the addition of the EMME features, suggesting that pre-trained model scores are providing some discriminatory signal orthogonal to one-hot encoding (**Fig. 2D**). Strikingly, while the validation balanced accuracy was high for both one-hot encoding and ESM-2 embeddings, only ESM2 embedding features resulted in 58.1% balanced accuracy on the test set, while one-hot encoding performed at 50% (expectation from randomly picking labels; **Fig. 2E**). While the improvement in performance is small, we reasoned that incorporation of EMME and ESM2 features does lead to improved generalizability, motivating a fuller use of the representations from these pre-trained neural network models for affinity prediction. All performance metrics are included in **Table S3.**

### Binding mutational landscape is smoothly connected and shows limited epistasis

A key question in molecular evolution, especially for relatively rapidly evolving viral systems, is the impact of epistasis on maintenance of function. If the joint functional impact of mutations is non-additive, the binding fitness landscape is expected to be rugged leading to potentially non-functional intermediates ^22,23^. On the other hand, if the impact of functional mutations is additive (lack of negative epistasis), we expect that evolutionary trajectories will smoothly sample functional diversity. Some previous studies^27,28^ analyzing deep mutational scanning data across diverse protein-protein interactions have found that impact of mutations is largely additive. As a result, one-hot encoded models trained on a limited amount of (single) mutational data are sufficient for predicting the function of combinatorial variants ^2,21,22,23^. Still others have found epistasis to be the dominant force for functional maintenance, making combinatorial effects “unpredictable” from the effect of single mutations^29–32^.

To illuminate the role of epistasis and the architecture of the evolutionary network at the RBD-ACE2 interface, we built logistic regression models with explicit one-body, two-body and three-body interaction features between one-hot encoded amino acids at positions sampled in our libraries ^21^. We chose LY005 and LY006 for this analysis as these libraries relatively densely sample combinatorial diversity at a small number of positions (5 and 6, respectively), and therefore provide good datasets for investigating the impact of amino acid interactions and epistasis on binding (**Fig. 3A,B,C; Table S4**). Feature sets were composed of one, two, and/or three-body interactions as binary variables following previous work ^21^. This partitioning allows us to estimate the degree of binding explained by epistatic interactions between pairs and triplets of mutations. To facilitate this analysis, we computed genetic scores - the sum of all main and epistatic effects contained in the corresponding variant - for each variant contained in the libraries (**Fig S5,S6**). In LY005, two- and three-body epistatic interactions only modestly contribute to activity, as performance accuracy only increased 4.9% over the one-body classifier. This is in contrast to the model trained on LY006, which exhibited a 9.7% performance increase in adding the two-body features to the one-body representations (**Figure 3A**). This implies that some two-body epistatic effects contribute significantly to binding in LY006. The overall discrimination between binders and non-binders was higher for the LY005 (**Figure 3B**) dataset compared to the LY006 dataset (**Figure 3C**). All performance metrics can be found in **Table S4.**

**Figure 3:**
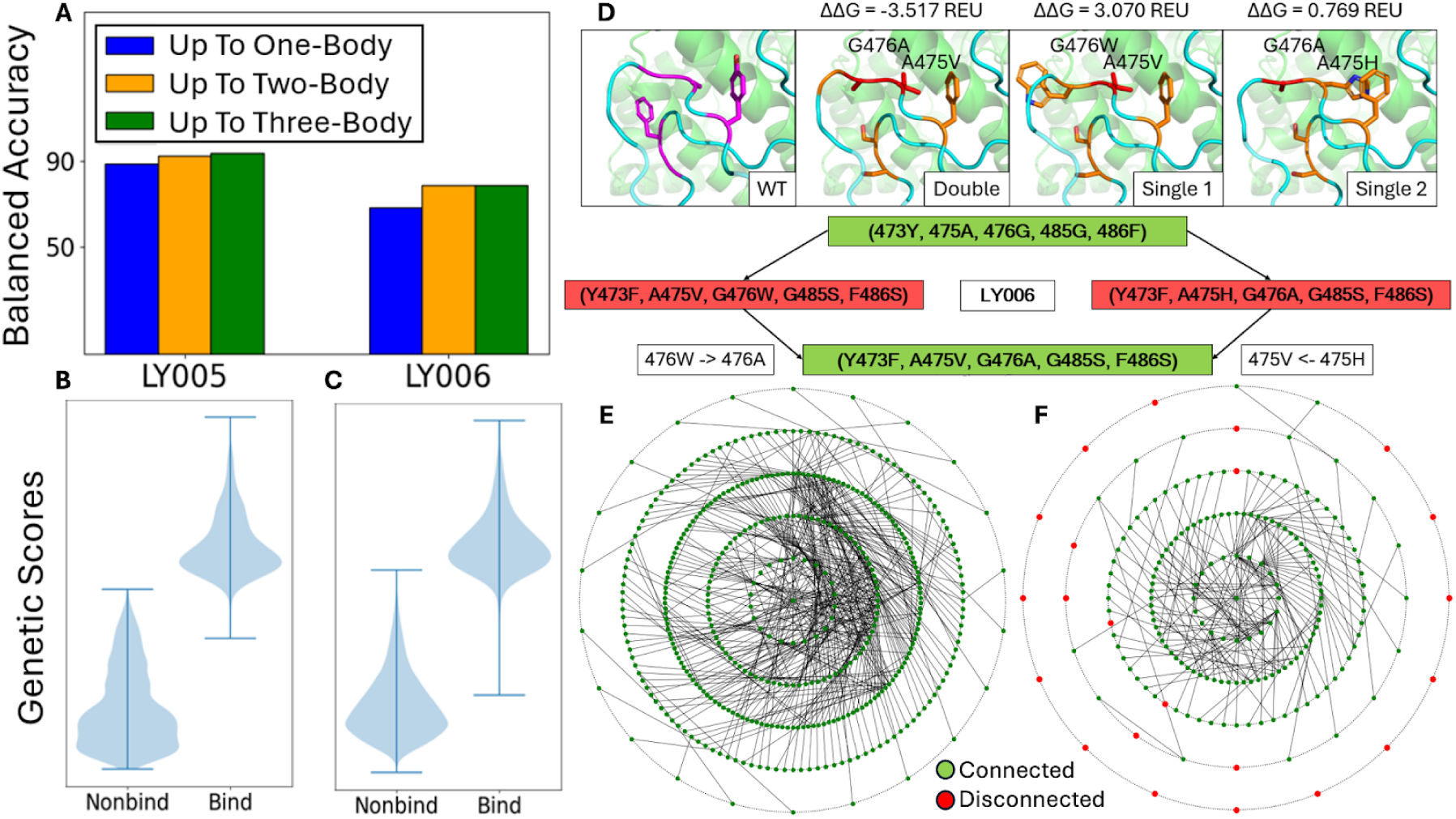
The S RBD mutational landscape is largely smoothly connected. **A:** Bar graph representing the best validation set balanced accuracies achieved by Logistic Regression using varying degrees of one-hot encoded sequence representations across 10 stratified training splits. **B, C:** Violin plots of the genetic score distributions for libraries LY005 (**B**) and LY006 (**C**) separated by binding class. **D:** Graphical representation of the sequences best showcasing pairwise positive epistatic interactions in library LY006. The positive epistasis exhibited between mutations A475V and G476A is depicted. Structural representations for the WT, the double mutant sample, and the two single mutant samples are shown at the top of the panel, alongside the ΔΔG of their total energies in REUs. Positions sampled in LY006 are shown in purple. Positions where mutations are installed are shown in orange. Mutations which correspond to either A475V or G476A are colored red. The RBD is shown in cyan, and ACE2 is colored green. We additionally depict the precise mutations installed in the double sample and its constituent single samples at the bottom of the panel. Samples shown in red are nonbinders, while samples shown in green are binders. The relevant WT amino acid identities are shown at the top of each diamond. The one-body samples are shown branching off of the WT, while the two-body sample is shown at the bottom of the diamond. The relevant point mutations installed to go from the one-body samples to the two-body sample are also included. **E, F:** Graphical representation of the sequence set accessibility analysis for libraries LY005 (**E**) and LY006 (**F**). The point at the center represents the WT sequence. Other points on the plot represent different designs in the library. Points on the Nth innermost circle represent sequences that are N mutations away from the WT. Only binding sequences are included in this representation. Green points depict sequences which can be connected back to the WT sequence by traversing a one-hamming-distance graph while maintaining binding activity for all nodes included in the walk. Red points represent sequences which cannot be connected back to the WT in this fashion. Edges represent the connections between sequences discovered while traversing the graph. Each edge connects two samples that are one mutation away from each other.

To investigate if the observed increase in performance accuracy upon inclusion of two-body terms in LY006 is indicative of epistasis or simply a reflection of adding more parameters to the (less well-performing) prediction model, we evaluated examples of predicted pairwise epistasis between positions in LY006. In our model, epistatic effects are computed relative to a “global average” of all variants in the library, not a single background sequence ^21^, making it difficult to directly interrogate isolated pairwise epistasis between two positions. Nonetheless, we used the sign and magnitude of logistic regression coefficients corresponding to all mutations in a given variant to identify predicted pairwise epistatic networks (**Supplementary Methods**). An example network between positions 475 and 476 (**Figure 3D**) involves the mutations A475V and G476A, which when made individually, are unfavorable for binding but together restore binding. Stability differences calculated using Rosetta are in agreement with the observed binding trends (**Figure 3D**): compared to the wild type, the single mutants have unfavorable ddGs of 0.8 and 3.1 REUs, indicating destabilization. The double mutant has a favorable ddG of -3.5 REUs, reflecting an increase in stability over the constituent single samples. Examining the energy contributions of different residues, we find that these differences arise from the solvation (fa_sol) and rotamer probability (fa_dun) terms for the residues 475 and 476. Thus, for this example, local interactions between these neighboring residues underlie the observed epistasis. The full energetic analysis is detailed in **Tables S5**.

To shed light on the implications of the observed epistasis on possible evolutionary trajectories, we investigated if each variant with known binding label can be reached via single mutational changes starting from the wild type RBD sequence. As expected based on the magnitude of the epistasis, all 410 variants which are binders in LY005 can be reached via single mutational trajectories (**Figure 3E**), whereas 22 of the 192 binding variants in LY006 (red points; **Figure 3F**) would require a non-functional intermediate to be reached via single mutation. While it is possible that the ruggedness observed in LY006 could be the result of lack of sampling of all possible variants in the experiments, the sampling is fairly dense and the average number of sampled neighbors in LY005 and LY006 are nearly identical (25.0 and 24.6, respectively), indicating that connectivity properties are not a reflection of sampling efficacy. Taken together, our investigation reveals that epistatic effects are minimal in the interface positions sampled in the LY005 library but may play a greater (but still limited) role in the residue positions sampled in LY006.

### New binders in diverse sequence space

Having observed that including scores from pre-trained DL models leads to improvements in prediction accuracies, we next investigated whether these models could be used to predictively generate new ACE2 binders. To that end, we used ProteinMPNN and ESM to sample at 12 positions directly at the ACE2 binding interface **(Fig 4A**) under the assumption of additivity (**Fig 3**). At each sampled position, we used each model (ESM or ProteinMPNN) to identify up to 5 top-scoring amino acids at each site. All identified single mutations were combinatorially enumerated and the generated sequences were scored according to the other model (ProteinMPNN or ESM, respectively) to identify those combinations that scored better than the Omicron variant. To ensure balanced sampling, we also randomly selected sequences from the combinatorially enumerated pools (see **Supplementary Methods** for more details). The library had a median of 7 mutations (range 1 - 8) per variant at 12 positions at the RBD-ACE2 interface (**Fig 4B**, **Fig 4C**). This combined library (LY010) was prepared as a yeast surface display library, sorted, sequenced, analyzed, and validated as described above (**Fig 1, Fig S3, Table S1**).

**Figure 4.**
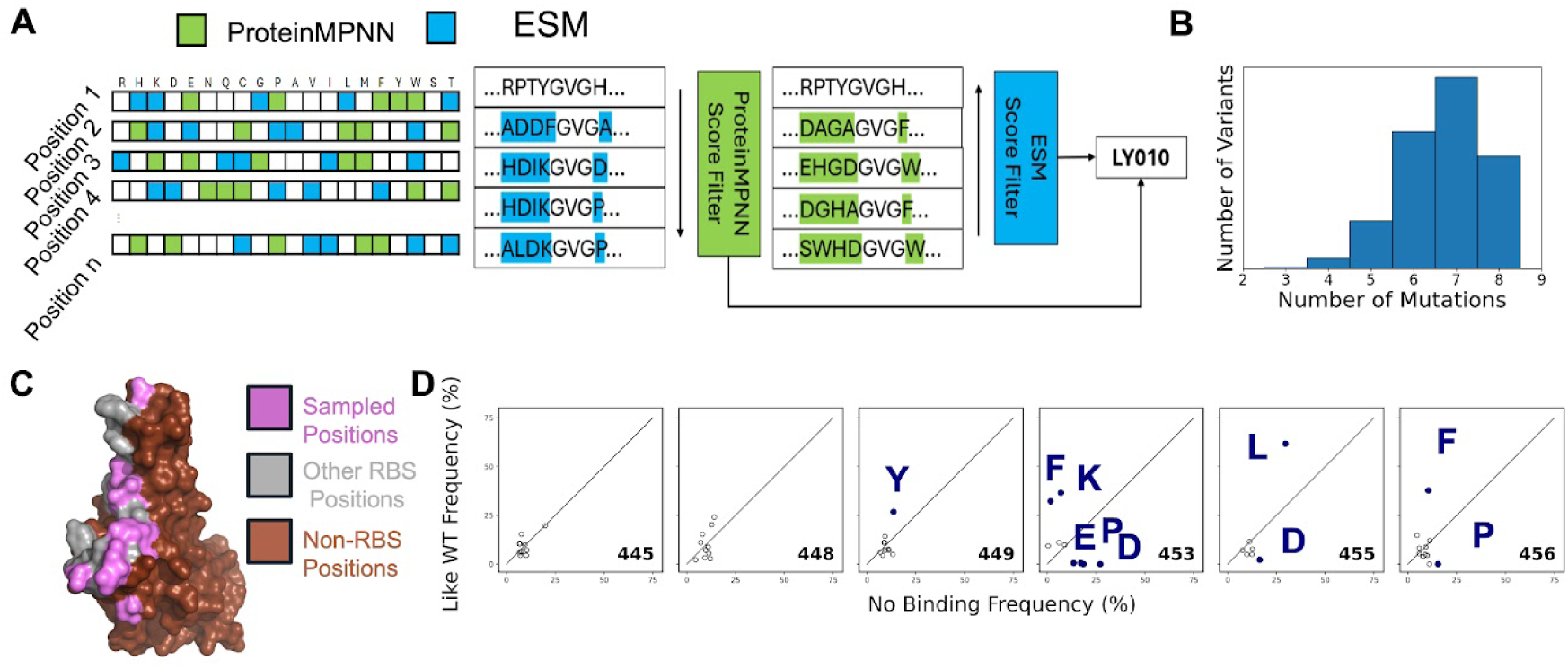
Computational design and screening of sequence diverse RBD variants. **A:** Graphic representing the design pipeline for library LY010. All possible point mutations at the target positions were first scored individually with ESM2 and ProteinMPNN. The top-scoring point mutations at each position were then assembled into full sequences combinatorially. The full sequences produced from each model were examined separately. The ESM2-suggested sequences were then scored and filtered by ProteinMPNN. Similarly, the ProteinMPNN-suggested sequences were scored and filtered by ESM2. The selected sequences from both approaches were then pooled together in LY010. **B:** Distribution of number of mutations for variants sampled in LY010. **C:** Structural representation of the sampled positions on the RBD. The structure is shown in surface view, with sampled positions colored pink. Other ACE2 interface residues are shown in gray, and the remainder of the structure shown in burnt red. The base structure corresponds to the Wuhan strain from PDB ID 6M0J. **D:** Per-position residue frequency differences between the ‘Like WT’ and ‘No Binding’ subsets from LY010. Sequences used in the plot had at least 100 sequence reads in the reference population and at least a 98% probability of belonging to their respective subset (“Like WT”: 183; “No Binding”: 1009 total sequences). Colored circles indicate residues where the difference in frequency was at least 12%.

We were able to recover 4,588 distinct sequences that could be assigned with confidence into the different binding categories. Of these, 1021 (22%) were ‘Like WT’, 1355 ‘Worse than WT’, and 2212 were non-binding (‘NB’) at 1 μM ACE2. The sequences that were classified as ‘Like WT’ were sequence diverse with a mean of 6.1 mutations (range 3-8). We selected two diverse sequences and validated that both had ACE2 binding affinity within 10-fold of the parental S RBS sequence (**Figure S3**). The sampling of the yeast library was particularly deep for cassette 2, allowing a thorough exploration of the functional landscape at positions [445, 448, 449, 453, 455, 456] of the RBD receptor binding site (RBS). Sequence profiling of these positions (**Fig 4D)** show considerable plasticity of the functional binding surface at these positions. Amino acid identities at positions 445, 448, and 449 were largely identical between the ‘No Binding’ and ‘Like WT’ subsets. At position 453, substitutions involving aromatic (F/Y) or large positively charged (H/K/R) were largely functional binders, whereas negatively charged (D/E) or proline substitutions were enriched in the non-binders subset (**Fig 4D**). At position 456 ACE2 binding preferences were for maintenance of its phenylalanine, whereas substitution to a proline was strongly disfavored for binding.

### Development of Graph Attention-Protein Language Model Network improves binding prediction accuracy

To investigate the ability of sequence and structure-based models for generalizable and accurate prediction of binding, we developed two additional neural network models relying on attention. In the first model, we applied a self-attention encoding layer to the ESM-2 embeddings of sequences in the dataset. In the second model, we processed Rosetta-generated structural models of dataset sequences into a graph representation as input into a graph attention module ^33^(**Figure 5A,B**). RBD residue positions at the RBD-ACE2 interface where sequence sampling was performed in any of the 4 libraries as well as their neighbors were considered as nodes of the graph (**Figure 5A**). Edges were included between spatially neighboring nodes. Edge features include C⍺ distance, sequence distance, and PyRosetta-computed interaction energy between the two residues. Node features include the average ESM embedding vector at the representative node, as well as the one-hot encoded representation of the amino acid at that position. Thus, every experimentally queried sequence is represented as a structure-based graph with geometric and energetic features (**Figure 5A**). Graph Attention^33^ was then applied to these representations and fed into a multi-layer perceptron for binding label prediction. Details of model architecture, training and validation are described in **Methods**.

**Figure 5:**
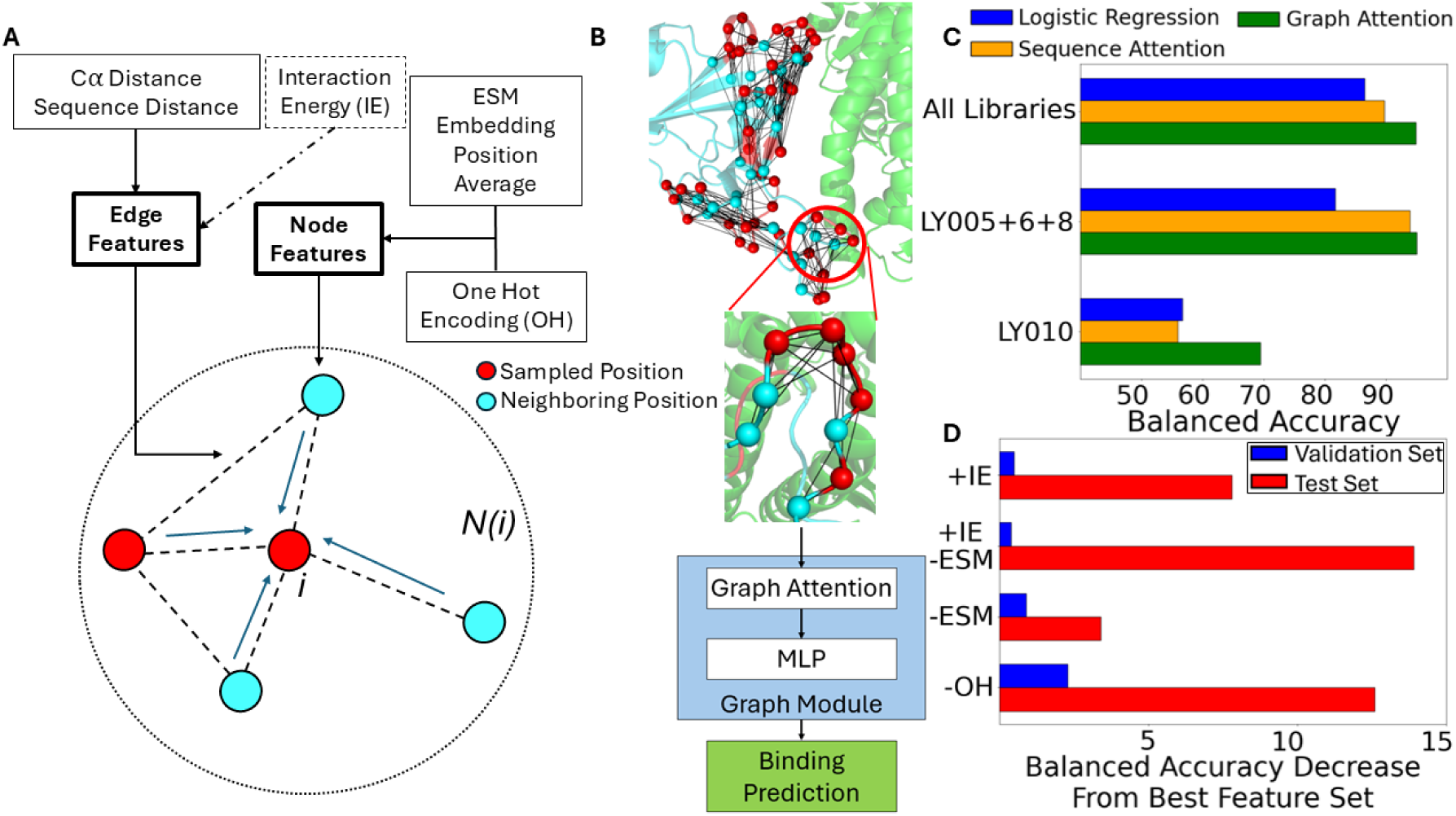
Graph attention improves generalizability at the RBD-ACE2 interface. **A:** Structure of the ACE2:RBD complex (PDB:6M0J) is used to compute a graph. Node features in this graph are the average of the ESM embedding at each residue position, the one-hot encoded amino acid identity, as well as an indicator variable reflecting whether the residue is located on the RBD or ACE2. Edge features include the C⍺ distance between the neighboring nodes, their sequence distance and (optionally) residue-residue pairwise interaction energy. Attention is computed between all neighboring nodes N(i) within 8Å for a given node i where sampling occurred. **B:** Encoded graph representations are passed into a graph attention convolution layer. The final representation is then passed through a multilayer perceptron (MLP) for final prediction readout. **C:** Prediction performance of the logistic regression, sequence attention, and graph attention models with different Training and Validation sets derived from combining various libraries. The first protocol consists of training and validating (10% data held out) on LY005, LY006, LY008, and LY010 (All Libraries, left). The second consists of Training and Validation (10% data held out) on LY005, LY006, and LY008 (LY005+6+8, middle), and testing on LY010 (LY010, right) **D:** Feature importance is reported as the decrease in balanced accuracy between the feature set perturbation and the best model on both the validation and test sets in the split protocol. Values greater than zero imply a decrease in performance upon addition or removal of the corresponding feature(s). (IE = pairwise Interaction Energy calculated using a decomposition of the Rosetta Energy Function; OH = One-Hot encoding; ESM = Model embeddings from the ESM-2 protein language model)

To investigate generalizability, sequences from libraries LY005, LY006, and LY008 were grouped into a single training set (90% training; 10% internal validation) and labels of all sequences in LY010 were predicted (test set). To obtain a ceiling for model performance, all libraries are used for training (with 10% internal validation sequences). All splits were selected randomly and stratified. Furthermore, all models used the same training and validation splits to ensure that their performances were comparable. The previously developed logistic regression model (**Fig 2**) was used as a baseline for comparison. When trained on libraries LY005+6+8 and tested on LY010, logistic regression attained 81.7% balanced accuracy on the internal validation set and significantly lower 56.7% balanced accuracy on the test set, indicating that this model fails to generalize to LY010 (**Figure 5C**). However, when trained on all libraries combined, the model achieved 86.5% balanced accuracy on the internal validation set, suggesting possible overfitting to training data. Similarly, the sequence-based attention model attained 93.9% (internal validation set LY005+6+8) but only 56.0% (test set) accuracy, and 89.7% balanced accuracy on the internal validation set when trained on the combined dataset. In contrast, the graph attention mechanism attained 94.5% (internal validation set), 69.5% (test set) and 94.8% (validation in combined dataset) balanced accuracies. The best performance was obtained for a model with sequence distance and C⍺ distance as edge features. The performance degraded considerably upon the addition of interaction energies computed from Rosetta as features, or the removal of either ESM embeddings or one-hot encoding features (**Figure 5D**). As both sequence and graph-based models use similar features derived from the RBD sequences, the performance improvement when using graph attention suggests that a more detailed and geometry-aware representation is necessary to improve generalizability. A full breakdown of all relevant performance metrics for the logistic regression, sequence attention, and graph attention models can be found in **Table 1**.

**Table 1:**
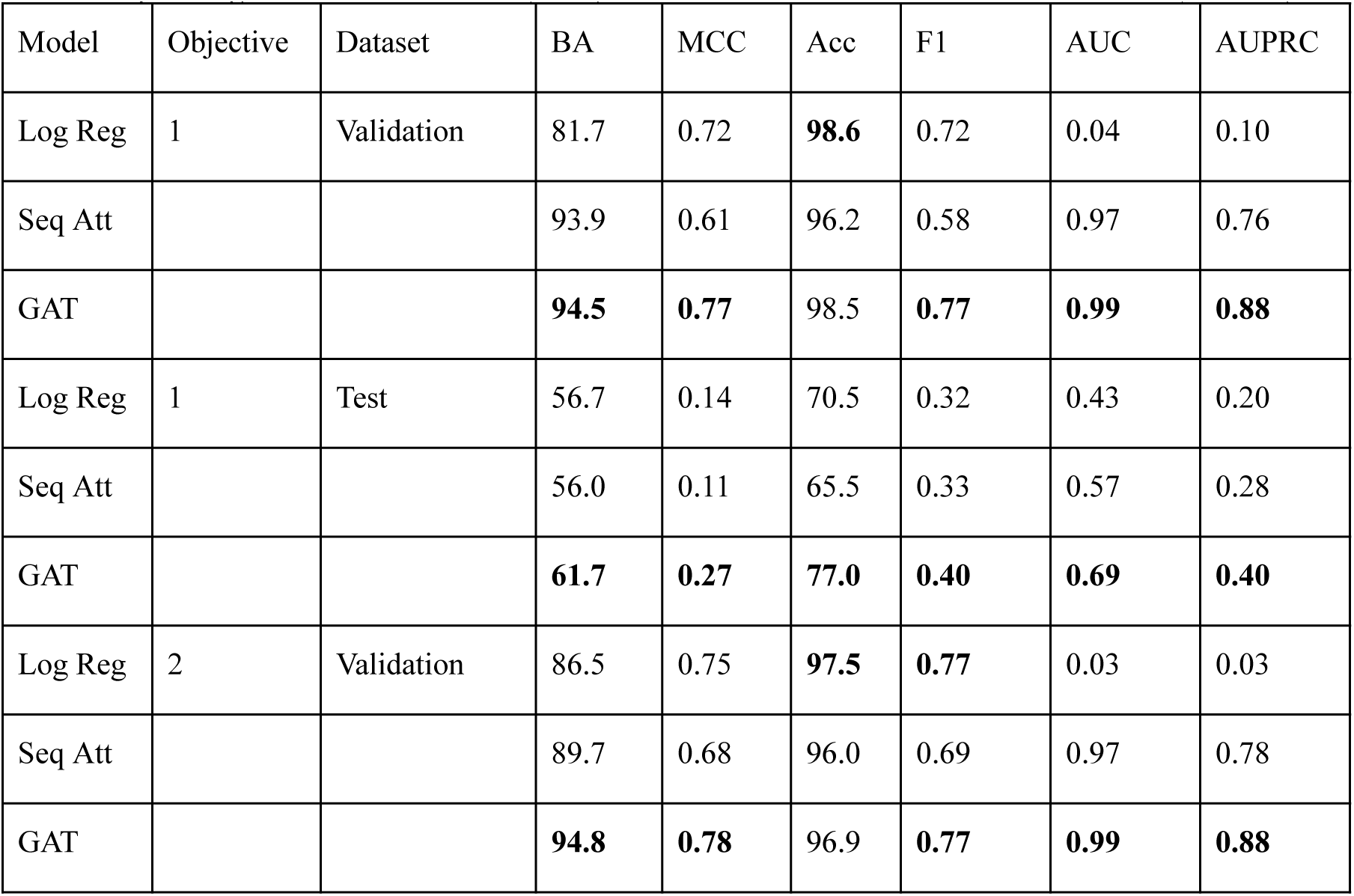
Various performance metrics attained with Logistic Regression (Log Reg), Sequence Attention (Seq Att), and Graph Attention (GAT). Objective refers to the two different training protocols employed, where models were either trained and validated on LY005+6+8 and tested on LY010 (Objective 1), or trained and validated on all libraries at once (Objective 2). Metrics included are Balanced Accuracy (BA), Matthews’ Correlation Coefficient (MCC), Overall Accuracy (Acc), F1-score (F1), Area Under the Receiver Operating Characteristic Curve (AUC), and Area Under the Precision Recall Curve (AUPRC).

## Discussion

It is well appreciated that plasticity is a key feature of the evolution of protein-protein interfaces, such that large numbers of mutations may accrue relative to the parental sequence during the course of evolution of organisms^8,34,35^. Facile access to variants of engineered proteins (such as antibodies and *de novo* designed protein binders) that maintain binding properties but are highly diverged in sequence compared to the parent sequence may also enable optimization of other therapeutically relevant properties such as immunogenicity, developability, and stability. However, identifying highly sequence distal variants given a parental variant is non-trivial. Experimentally, since random mutations are on average deleterious, randomly sampling variants distant from parental sequences result in exceedingly low probabilities of functional sequences. Furthermore, due to the context dependence of single mutations, predicting functional effects in sequences far from the parental sequence using single (deep) mutational scans on the parental background is also challenging. Computationally, while physics-based approaches such as molecular dynamics simulations with free energy based calculations can provide accurate estimates of the changes in binding free energy upon multiple mutations, these are prohibitively expensive. On the other hand, relatively more inexpensive methods, e.g., FoldX^36^ and Rosetta^37^, are limited in their ability to predict binding affinity changes beyond a handful of amino acid substitutions due to the many approximations underlying these approaches. We reasoned that deep learning (DL) models of proteins such as ESM and MPNN, which have been trained on broad swathes of the protein universe to develop general representations of viable sequence-structure relationships, may enable more efficient navigation of remote, functional sequence space to yield balanced datasets containing larger numbers of binders. Indeed, we found that the MPNN- and ESM- sampled library LY010 contained a significant number of Like WT binders (22.3%). Compared to the focused combinatorial libraries LY005 (5.3% Like-WT binders) and LY006 (1.0%), the functional percentage of variants in LY010 is considerably higher despite many of the mutated positions being shared across multiple libraries. These results demonstrate that the incorporation of generalizable model-derived features improves sampling over alternate methods (**Table S7**). Thus, using DL models improved the breadth and depth of sampling of functional binding proteins at one side of an interface, which is more extensive compared to previous efforts on S RBD^7,10,11^ and other protein interactions^38,39^. In total, we experimentally evaluated 43,000 sequence variants that contain up to 26 substitutions compared to the parental chimeric Omicron RBD sequence. We observed functional RBD variants with up to 23 mutations from the parental sequence, and binders with up to 8 mutations directly at the receptor binding site which maintains parental ACE2 binding. Our results point to the allowable, functional sequence space at the RBD being immensely large, consistent with the continued accrual of mutations at the RBD in circulating viruses.

While DL models of proteins aid in increasing sampling efficiency in functional sequence space, their ability to accurately predict function remains limited. Incorporation of DL model representations as features (e.g., EMME features in **Fig. 2** or ESM embeddings in **Fig. 5**) in classifiers leads to slight improvements when combined with the baseline one-hot encoding, but fails to effectively generalize to test sets. The reliance on one-hot encoding limits model generalizability as this representation implicitly captures the neighborhood of the protein around a specific residue position which results in a single amino acid preference. DL model embeddings and the architectures used to convolve these embeddings to predict function - binding in our case - clearly fall short in effectively capturing these neighborhoods. One avenue for improvement which may abrogate the need for one-hot encoding and improve generalizability may be training models using binding measurements with a diverse set of binding interaction proteins as starting points^40^. Training the prediction models on a wide variety of environments across a structurally diverse set of proteins may allow learning high accuracy representations of sequence-structure-function relationships underlying protein-protein interaction specificity.

Many different computational methods have been employed to predict the outcomes of protein-protein interactions, ranging from standard machine learning algorithms to more complex deep learning architectures. Baseline approaches have often utilized models such as Random Forest, Support Vector Classifiers, and Logistic Regression, typically paired with one-hot encoded sequence representations^11^. While these can effectively capture site-wise amino acid preferences within a given dataset^2^, our work confirms they fail to generalize to highly sequence-diverse libraries, especially if residue positions not seen in the training dataset are present in the test set (**Fig. 2**). Neural network models that rely on sequence-based features alone, such as applying attention to protein language model (PLM) embeddings, likewise show limited generalizability^41–43^. Our graph attention network (GAN-PLM) architecture was developed to overcome these limitations by combining the advantages of both structure- and sequence-based methods. This approach naturally incorporates protein structure by representing the interface as a graph, using structural models to define nodes (residues) and edges (spatial proximity). This graph framework allows the model to explicitly consider interface geometry, with edge features like Cα distance, while integrating rich sequence information—including PLM embeddings and one-hot encodings—as node features. By unifying these representations, our graph attention model explicitly captures residue environments and sequence dependencies, achieving substantially higher balanced accuracy and generalizability when predicting function across diverse, unseen sequence libraries (**Fig. 5**). We found that explicitly considering residue-residue pairwise energies as features in the graph does not aid prediction accuracy unlike our previous work in predicting protease-peptide cleavage specificity^44^. This difference may arise due to the greater flexibility and diversity of environments at the RBD:ACE2 interface compared to the very restricted and exacting orientation requirements for enzyme:substrate complexes, as energies are highly sensitive to the small perturbations in structure. Nevertheless, consideration of protein geometry in our GAN-PLM contributes significantly to the prediction accuracy as indicated by the worse performance of the sequence-attention-based model which has all “node”-based features in GAN-PLM but no protein geometry information (**Fig. 5**).

There are limitations in our construction of deep mutational scanning datasets that impact interpretability and generalizability of the data. First, our binding analysis is qualitative and is pegged to the dissociation constant of the parental RBD sequence used in the sorts. While in principle quantitative dissociation constants can be measured using yeast display coupled to deep sequencing ^3,45–47^, such measurements require a higher percentage of ‘like WT’ binders than found in the initial libraries tested (LY005, LY006, LY008). Second, our qualitative binning method found 93% agreement between biological replicate libraries and 87% agreement between population measurements and titrations on individual variants. These values set an upper bound on agreement between models and the experimental dataset. While current test set accuracies (e.g., <70% in Fig. 5C) are far below the experimental levels, this issue will gain salience as prediction methods improve. Third, we performed all measurements in the context of the same parental sequence. Full generalizability for a given protein interface likely requires binding measurements using many different binding interaction proteins as starting points^40^. Fourth, we did not include insertions or deletions in our experiments. Since indels are accessible by deep mutational scanning experiments^48,49^, inclusion of such sequence variants are an avenue for further exploration. Fifth, we did not incorporate mutations away from the interface (in the protein core, for example), which could alter the protein fold or rigid body placements of the ACE2 contacting residues and, thus, their preferences^50^. Sixth, our Golden Gate assembly strategy segmented the receptor binding site over two cassettes. All variants tested had mutations in one cassette only, which limits our experimental sampling over the entire protein. Advances in gene synthesis, particularly for larger length genes, would help better sampling across the entire protein domain. Overall, these limitations result in our datasets being a considerable lower bound on the sequence tolerance at the RBD protein interface.

There is continued debate about the extent of epistasis present in natural proteins and the appropriate reference for determining epistasis. As sequences drift farther away from the parent sequence, it is expected that epistatic effects play a greater role as mutational environments and/or protein structure changes significantly. In our experiments, we observed modest but detectable epistasis especially in the larger LY006 library; our experimental and ML approach allows delineation of these global epistatic effects. Better understanding how epistasis occurs allows effective prediction of multiple mutation effects and enumeration of the tolerated sequence space^51^. Note that our definition of fitness used here - maintenance of the RBD fold and binding to ACE2 - understates the true epistatic relationships present in the SARS-CoV-2 Spike protein *in vivo*. For example S RBD mutations K417N & N501Y observe an epistatic sign effect, where K417N contributed to immune evasion at the expense of binding affinity. N501Y recovered binding affinity in this background ^52,53^. Our measurements are also made in the context of a single RBD displayed on the surface of yeast, and conformational equilibria associated with the full-length S trimer may modulate some of these mutational effects.

In summary, we developed a combined computational-experimental approach to illuminate large swathes of functional sequence diversity on one side of a protein-protein interface. Sampling informed by the DL models ESM and ProteinMPNN improved the efficiency of identification of variants that differ by >20 amino acid substitutions compared to the parent sequence and a graph neural network with protein language model embeddings improved binding prediction accuracy on unseen sequences. Continued development of experimental and computational methods and application of our approach to a larger diversity of protein-protein interfaces should make the accurate and exhaustive enumeration of sequence diversity compatible with protein-protein interfaces in a zero-shot manner more feasible, and enable further applications in therapeutic protein design and synthetic biology.

## Materials and Methods

### Sequence design of mutagenic libraries

The parental sequence, denoted ‘WT’, for all mutagenesis experiments was a chimeric SARS-CoV-2 S RBD (333-541) Wuhan-1 with common mutations found on Omicron sub-variants (K417N, S477N, E484A, Q498R, N501Y and Y505H) ^19^. Sequence design strategies are described below in the Computational modeling section. All the sequences are provided in **Supplementary Data.**

### Preparation of the mutagenic libraries

Libraries were constructed exactly as described in Daffern et al ^19^ using degenerate oligonucleotides (LY005 and LY006) or custom oligo pools (LY008, LY0010) sourced from Agilent. The constructed libraries were prepped using ZymoPURE II plasmid midiprep kit to extract the plasmid. Yeast transformation was performed exactly as described ^54^ by transforming 5 µg of DNA into chemically competent *S. cerevisiae* EBY100 ^55^. Serial dilutions were plated on SDCAA agar (20g/L dextrose, 6.7g/L Difco yeast nitrogen base, 5g/L Bacto casamino acids, 5.4g/L Na_2_HPO_4_, and 8.56g/L NaH_2_PO_4_·H_2_O supplemented with 15 g/L agar) plated and incubated for 3 days at 30°C to calculate the efficiency of the transformation. In parallel, the cells were grown in SDCAA liquid media (20g/L dextrose, 6.7g/L Difco yeast nitrogen base, 5g/L Bacto casamino acids, 5.4g/L Na_2_HPO_4_, and 8.56g/L NaH_2_PO_4_·H_2_O) with 50 𝜇g/mL Kanamycin and 1x PenStrep for 3 days at 30°C and 300 rpm to saturation for 30h. For medium term storage at -80°C, yeast stocks were prepared at OD_600_ = 1 and at OD_600_ = 10 for screening and sorting respectively as in Whitehead et al. ^56^. Each library contained between 99.1% and 100% of the desired variants, as confirmed by Illumina short read sequencing (full details given in **Supplementary Data**).

### Screening and sorting libraries

For cell surface screening the cell stocks were grown in SDCAA at OD_600_ = 0.5 overnight at 30°C and 300rpm. The next day, the cells were induced in SGDCAA (18g/L **g**alactose, 2g/L **d**extrose, 6.7g/L Difco yeast nitrogen base, 5g/L Bacto casamino acids, 5.4g/L Na_2_HPO_4_, and 8.56g/L NaH_2_PO_4_·H_2_O) at OD_600_ = 1 at 22°C and 300 rpm for 22h with 50 𝜇g/mL Kanamycin and 1x PenStrep for 30h. For sorting the libraries, 2% of the ‘WT’ was spiked in on the SGDCAA inducing culture. The displaying cells were washed with PBSF (PBS containing 1g/l BSA) and stored on ice until used.

For sorting of the libraries, 3x10^7^ cells were labeled at indicated concentrations of ACE2 (University of Washington, Institute for Protein Design, human ACE2-Fc, Lot #20200706) in PBSF (1nM, 10nM, 100nM and 1000nM) for 1h at 22°C with shaking. The cells were centrifuged and washed with 1000 𝜇L PBSF. Each reaction was split into four Eppendorf tubes, except for the LY010 library and individually labeled with 2.4 μL anti-c-myc-FITC (Miltenyi Biotec), 1 μL Goat anti-Human IgG Fc PE conjugate (Invitrogen Catalog # 12-4998-82) and 196.6 μL PBSF for 10 min at 4°C. Cells were then centrifuged, washed with PBSF. The pellet was resuspended in 1000 𝜇L and read on a flow cytometer to measure binding of the ACE2. The gates used for sorting were configured using cells not labeled with ACE2 and are shown in **Figure S1**. The gates set are the following: an FSC/SSC^+^ gate for isolation of yeast cells, FSC-H/FSC-A gate to discriminate single cells, a FSC-A/FITC+ gate selects the cells displaying the RBD on their surface and from this last gate, everything above background noise by a PE^+^/FITC^+^ is collected.

Cells were collected at approx. 100-fold of the theoretical library coverage (1.0x10^6^ for LY005, LY006, LY010, and 5.0x10^5^ for LY008 and LY010). A reference for each library was also collected as the cells that display in the FSC-A/FITC+ gate at approx. 500-fold of the theoretical library coverage (5.0x10^6^ for LY005, LY006 and 3.0x10^6^ for LY008 and LY010). The collected cells were spun down and recovered in SDCAA with 50 𝜇g/mL Kanamycin and 1x PenStrep for 30h. Replicates of the LY008 library were sorted on different days using different yeast stocks of the same Golden Gate reaction.

### Yeast surface titrations of individual variants

Random library members from LY005, LY006, LY008, and LY010 were chosen for individual titrations. Additional members from each of these libraries were also deliberately chosen to include variants predicted to bind like WT.

Libraries were separately plated on SDCAA Agar plates and grown at 30°C for 2 days. Individual colonies were selected and grown in SDCAA with 50 𝜇g/mL Kanamycin and 1x PenStrep at 30°C and 300 rpm for 30h. The cells were induced in SGDCAA with 50 𝜇g/mL Kanamycin and 1x PenStrep at 22°C and 300 rpm for 22h. For LY0005, ten variants were chosen randomly, and one was assembled from gBlocks (IDT) using Golden Gate assembly and transformed into yeast (2). For LY006, ten variants were chosen randomly and one was constructed via Golden Gate assembly. For LY008, seven variants were chosen randomly and two were constructed via Golden gate. For LY0010, six variants were chosen randomly, and two were assembled via Golden Gate assembly. Titrations were performed as described by Chao et al. ^57^ using a concentration range of 1 pM to 1 µM ACE2 and with an incubation time of 4h at 22°C in a shaking incubator. Variants LY005v12, LY006v11, LY008v11, and LY008v12 were titrated with ACE2 supplied by University of Washington, Institute for Protein Design (human ACE2-Fc, Lot #20200312). For LY010, variants 11-14 were titrated with ab273687 ACE2 (Abcam). All other variants were titrated with the same ACE2 as was used for sorting (University of Washington, Institute for Protein Design, human ACE2-Fc, Lot #20200706).

### Polyspecificity assay

Lysozyme was biotinylated with EZ-link NHS-Biotin following manufacturer’s instructions. 1x10^5^ yeast cells were labeled with 250nM, 1µM and 5µM biotinylated lysozyme (Sigma L6876) for 30 min at 22°C and shaking. The cells were centrifuged and washed with 200𝜇L PBSF. The cells were secondary labeled with 0.6 𝜇L anti-c-myc-FITC (Miltenyi Biotec), 0.25 𝜇L SAPE (ThermoFisher S688) and 49.15𝜇L PBSF for 10min at 4°C. The cells were washed and resuspended in 100𝜇L PBSF. 25,000 cells were screened using a flow cytometer for each labeling concentration.

### Deep sequencing preparation

The DNA was prepared for deep sequencing following the “Method B” protocol from Kowalsky et al ^58^. The amplicon was amplified as described in Daffern et al. ^19^. Samples were then further purified using Agencourt Ampure XP beads (Beckman Coulter), quantified using PicoGreen (ThermoFisher), pooled, and sequenced on an Illumina MiSeq using 2 x 250 bp paired-end reads at Rush Genomics and Microbiome Core Facility (Rush University, Chicago, IL).

### Sequencing analysis and variant labeling

For the analysis of the RBD library deep sequencing data, sequences were merged using an in-house merging code, essentially as previously described by Haas et al. ^59^. Reads were filtered to remove any variants that were not expected in the designed libraries, except for the 1 nM selected population for LY0010. This population had aberrantly large reads of WT sequences likely due to cross-contamination during PCR. This population was re-sequenced separately by preparing amplicons from frozen yeast stocks, merging reads using Flash ^60^, filtering to remove reads with quality scores <30, and removing any variants that were not expected in the designed libraries.

The observed probability of sorting each variant i at a labeling concentration j (p_ij_) was determined using the following equation:

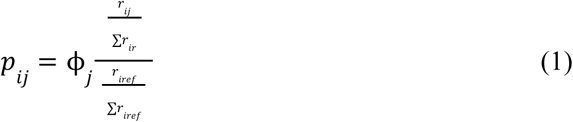

Here ϕ_𝑗_ is the sorted fraction of cells at each labeling concentration j, relative to the reference control, r_ij_ is the number of observed reads of variant i at labeling concentration j, and r_iref_ is the number of observed reads of variant i in the reference population. For the 1 nM selected population in LY010, Σ*r_ir_* was corrected to account for contamination of this population with wild-type variants. Under the assumption that the true frequency of wild-type in the original population matches the frequency in the re-sequenced population, and that the originally sequenced variant count is correct, the corrected Σ*r_ir_* was calculated. For each variant in this population except the wild-type, 𝑟_𝑖,𝑗=1 𝑛𝑀_ was set equal to the sum of read counts from the initial and repeat sequencing data. The 1 nM wild-type probability was calculated based on variant and total counts from the resequencing data only.

A score for the likelihood of variant i belonging in one of five classification bins k (‘binds with affinity similar to WT’, ’10-fold worse’, 100-fold worse’, ‘1000-fold worse’, ‘no binding observed’) was determined from the following equation:

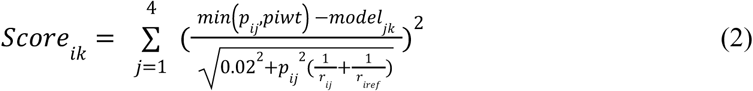

Where the vector for the model predictions are drawn from the following table:

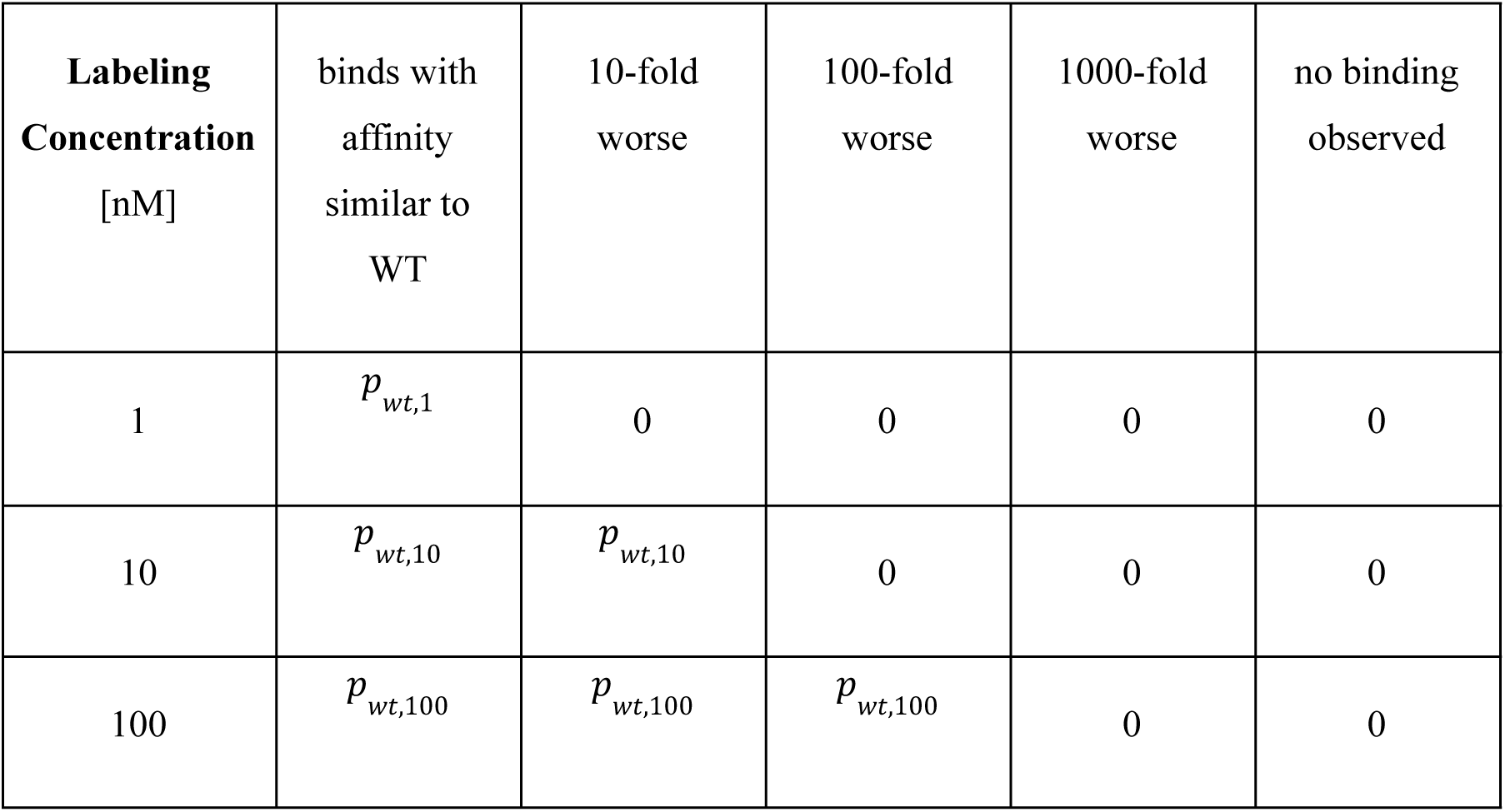
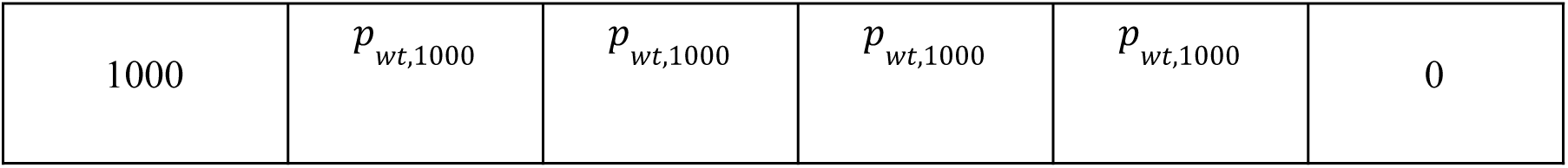

The normalized probability of each variant i falling into a classification bin k (PROB_ik_) is computed using an inverse F-statistic test with 3 degrees of freedom (df) and 1 parameter:

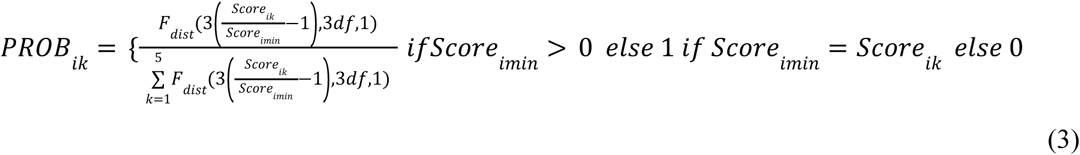

The dataset was reduced to three bins: “like WT”, “worse than WT” and “no binding”. From the initial five bins, “like WT”, “100-fold worse than WT” and “no binding” stayed the same. For the middle bins (“10-fold” and “1000-fold worse than WT”), the surrounding probabilities were taken into consideration and the variant was classified according to the higher probability of the neighboring bins.

### Computational sequence selection for LY005, LY006, and LY008

We designed four libraries with different approaches. Libraries LY005 and LY006 were designed based on the structural analysis of the RBD-ACE2 interaction and were made with degenerate codons with mutations to Class 2 (LY005) or Class 3 (LY006) epitopes. To limit the number of mutations in these libraries, we used a reduced amino acid alphabet ^17^. Residue selections were made based on their prevalence in previous investigations into the impact of point mutations on the RBD:ACE2 interface. The sampled positions in both LY005 and LY006, as well as the allowed residues at each position, are highlighted in **Table S6**. The native amino acids are included as the first entry of each position list.

We then used ProteinMPNN to generate sequences for each cassette separately. Several different design approaches were implemented, including restricting sampling to observed mutations mined from the GISAID database^18^. A full detailing of the computational design protocol can be found in **Supplementary Methods**.

Structural models were made for all designs by grafting the MPNN mutations onto the native structure (PDB ID 6M0J) using PyRosetta FastRelax^61^. The 𝚫𝚫Gs for binding and folding were then computed using both PyRosetta and the established Cartesian 𝚫𝚫G protocol^62^. Designs were filtered down according to their computed energies, yielding the final sequences for library LY008.

### Baseline Assessment

Support Vector Classifier, RandomForest, K-Nearest Neighbors, Naive Bayes, and Logistic Regression were implemented using the Scikit-learn package in Python^63^. All models were trained using stratified 10-fold cross-validation. A full breakdown of the best attained performances can be found in **Table S2.**

### Analysis of epistasis

To compute the epistatic effects of pairs and triplets of states, we first extracted the coefficients from the full model and matched them to their corresponding mutation sets. These model parameters were then used to compute the genetic scores of all variants. The genetic score of a given sample is defined as the sum of all main and epistatic effects contained in the combination of sequence states present in the corresponding variant. Therefore, an increase in a sample’s genetic score is linearly correlated with its probability of binding Like WT. Once determined, these genetic scores enabled us to calculate the main and epistatic effects of all states in both libraries LY005 and LY006 (**Fig S6, S7**). Using the computed main and epistatic effects for all states, pairs of states, and triplets of states, we identified promising examples of positive pairwise epistasis in both libraries LY005 and LY006. A number of constraints are imposed to find the clearest examples of epistasis, including a consistent mutational background between the double and single samples, as well as ensuring that only the double samples bind ACE2. The full procedure is detailed in **Supplementary Methods**.

### Energetic Validation

We closely examined the samples that logistic regression identified as exhibiting pairwise positive epistasis to verify our epistatic model. We selected double beta design (Y473F,A475V,G476A,G485S,F486S) from LY006, featuring target state (A475V,G476A), as this sequence had the largest difference in binding probability between itself and its constituent single beta samples. These single designs were (Y473F,A475V,G476W,G485S,F486S) and (Y473F,A475H,G476A,G485S,F486S).

We next examined the PyRosetta-generated structural models for the double sample and both single samples to determine if energetic measurements could corroborate our epistatic analysis. To this end, we computed the total energy of each sample across the whole RBD. These scores were subtracted from the native to obtain ΔΔG values for the single and double samples.

The energetics at each sampled position in LY006 were next examined to determine the precise cause of the total score discrepancy. Position selectors were initialized separately for positions 456, 473, 475, 476, 485, and 486. The total score was decomposed into its constituent energy terms, so each component could be investigated individually. The average of the differences between the double sample and the two single samples for each score term was calculated, and used to find the biggest contributors. The 5 largest discrepancies where the double sample is significantly more favorable than the single samples, as indicated by a lower energetic term difference, are shown in **Table S5**. The total_score score term was excluded for this analysis.

### Selection of sequences using ESM and ProteinMPNN

The LY010 library was constructed using multiple pre-trained models to suggest functional areas in sequence space to be sampled from. To reduce the size of the search space, only particular positions were sampled for each cassette, as detailed in **Supplementary Methods**.

To determine which mutations should be installed, three separate scoring approaches were devised. For the first approach, point mutations were installed individually across the whole cassette, yielding (19*N)+1 total sequences. Each point mutation was scored with ESM2 relative to the chimeric Omicron parental sequence. The top 5 scoring mutations at each position were retrieved, and designs were assembled combinatorially and exhaustively. The full sequences were then scored using ProteinMPNN^6^. The differences between design scores and the omicron variant score were computed and used as a filter. t-SNE dimensionality reduction ^20^ was performed on the passing sequence set, which was further filtered based on a minimum 2-dimensional distance threshold to the rest of the set.

The second approach is the reverse of the first method. The point mutations were scored using ProteinMPNN to select the best 5 at each sampled position. These mutations were then sampled combinatorially to generate full sequences, which were scored by ESM2 relative to omicron. Sequences with negative ESM2 scores were discarded. Dimensionality reduction was conducted identically to the first approach, and designs were filtered by minimum geometric distance.

The third approach combines approaches one and two. The best point mutations at each position as determined by both ProteinMPNN and ESM2 in the first two approaches were combined and sampled together combinatorially for each set of positions. Dimensionality reduction and distance filtering was performed as previously described. The full set of ProteinMPNN and ESM2-suggested sequences are referred to as library LY010.

### Development of Attention-Based Models

We employed an attention-based model to process sequences encoded by ESM2. The model first took as input the 2560-dimensional length-averaged vector for each sequence, then passed it through a self-attention encoder prior to prediction readout. We encoded our designs into a graph representation for input into a graph attention-based model. To begin graph construction, all positions along the RBD which were sampled anywhere in the four libraries were retrieved.

Positions on ACE2 and RBD which fell within an 8Å C⍺ distance cutoff (Based on PDB structure 6M0J) to any sampled position were also included. All identified positions served as nodes in the RBD graph. Edges were added between nodes under the 8Å C⍺ distance threshold. Edges were bidirectional, and encoded with the C⍺ distance, sequence distance, and PyRosetta-computed interaction energy between the two neighboring nodes. Inside of the nodes, we also included the average of the ESM embedding vector at the representative position, as well as the one-hot encoded representation of the amino acid at that position. A position indicator was also added, which specified whether the node was on the RBD or ACE2 (**Figure 3A**). The encoded graph representations were then used as input into our graph attention model. All relevant architectural details and hyperparameter selections can be found in **Supplementary Methods.**

## Material availability

All the sequences have been deposited in SRA (SAMN36730863-SAMN36730913; SAMN41763486-SAMN41263490; SAMN52945872). Processed deep sequencing datasets are available at 10.5281/zenodo.17485853.

## Code availability

The code used to process raw sequencing data is available via GitHub at https://github.com/WhiteheadGroup/Graph-Attention-SARS-RBD.git

The code used for training the graph and sequence attention models, as well as performing the epistasis and neighborhood analyses, is available via GitHub at https://github.com/JonathanEAsh/GraphAttentionRBD.

The PyRosetta-generated structures for all RBD variants can be downloaded alongside the precomputed ESM2-embedding averages from our Zenodo repository [here](https://zenodo.org/records/15548841?token=eyJhbGciOiJIUzUxMiJ9.eyJpZCI6ImFlOTQwNjkwLTcwNTctNDQwMS1hYjBjLTI1MjMwMzBlODEyOSIsImRhdGEiOnt9LCJyYW5kb20iOiI0M2NjMzNmZjdjNzUxYTVmMGZhMGE2YjQ0YWUwNDI2MCJ9.0uT8f_yjJ-w9nXE6oh5hFM-IjnF2zGoXZsG0lpqur1ocCBmWND2owyXILlff2YmLbQosUOooCMbpvrZZawXz6Q).

## Acknowledgments

The authors would like to thank Neil King for the kind gift of Fc-ACE2. This work was supported by the National Institute Of Allergy And Infectious Diseases of the National Institutes of Health (Award Numbers 5R01AI141452-05 to T.A.W.; Award R21AI174157 to T.A.W. and S.D.K.), the NIH/CU Molecular Biophysics Program and NIH Biophysics Training Grant T32 GM145437 (participant support, S.P.K.), and the NSF RAMP Post-Bac Training Program Award #2216011 (participant support, C.N.D.).

## Author Contributions

Designed research: JA, IMF, SPK, TAW, SDK; Performed research: JA, IMF, SPK, CND; Wrote the paper: JA, IMF, TAW, SDK; Supervised research: TAW, SK

## Competing Interest Statement

TAW is a consultant for Inari Ag and serves on the scientific advisory board for Metaphore Biotechnologies and Alta Tech.

