## Supplementary material for "Graph attention with structural features improves the generalizability of identifying functional sequences at a protein interface": Suplemental Information

### Supporting Information for
Graph attention with energy features improves the generalizability of identifying functional sequences at a protein interface

Ash, J.[2♀], Francino-Urdaniz, I. M.[1♀], Kells, S. P.[1], Davis C. N.[1], Whitehead, T. A.[1,*], Khare, S.D.[2,*].

[1]Department of Chemical and Biological Engineering, University of Colorado, Boulder, CO, 80303, United States of America

[2]Department of Chemistry & Chemical Biology, Rutgers The State University of New Jersey, 123 Bevier Rd, Piscataway, NJ 08854, United States of America

*Whitehead, T. A., Khare, S. D.

♀These authors contributed equally

**This PDF file includes:**

Figures S1 to S7
Tables S1 to S8
Supplementary Methods
Legends for Datasets S1 to S5
SI References

**Other supporting materials for this manuscript include the following:**
Datasets S1 to S5
Supplementary Data S1: Primers, plasmids, sequences, transformation efficiencies

#### Supplementary Figures

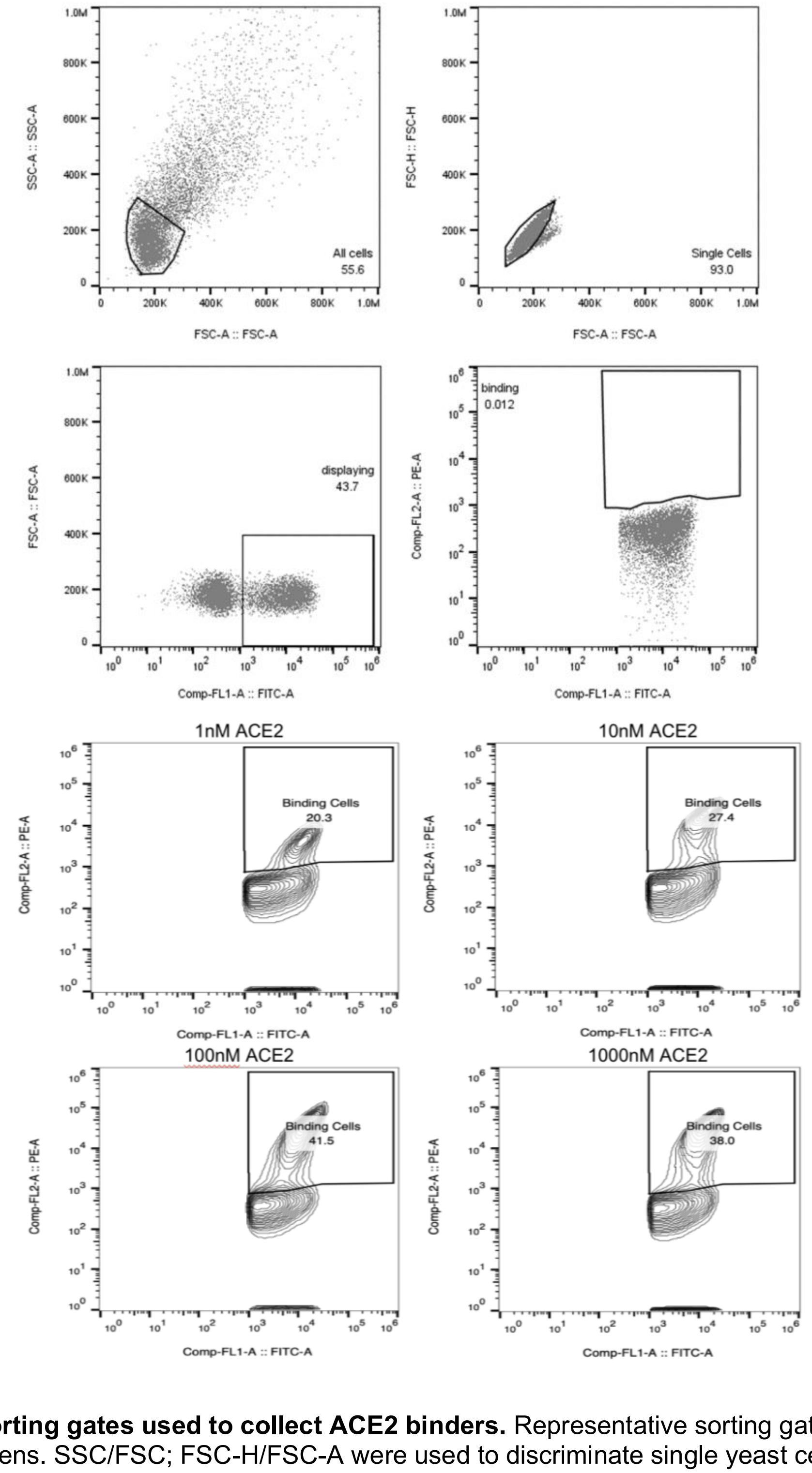


**Fig. S1. FACS sorting gates used to collect ACE2 binders.** Representative sorting gates used for all library FACS screens. SSC/FSC; FSC-H/FSC-A were used to discriminate single yeast cells; FSC-A/FITC+ to select cells displaying the RBD on their surface; and $SAPE^{+}/FITC^{+}$ to identify mutants that bind to ACE2 above the cells displaying the RBD and noise.

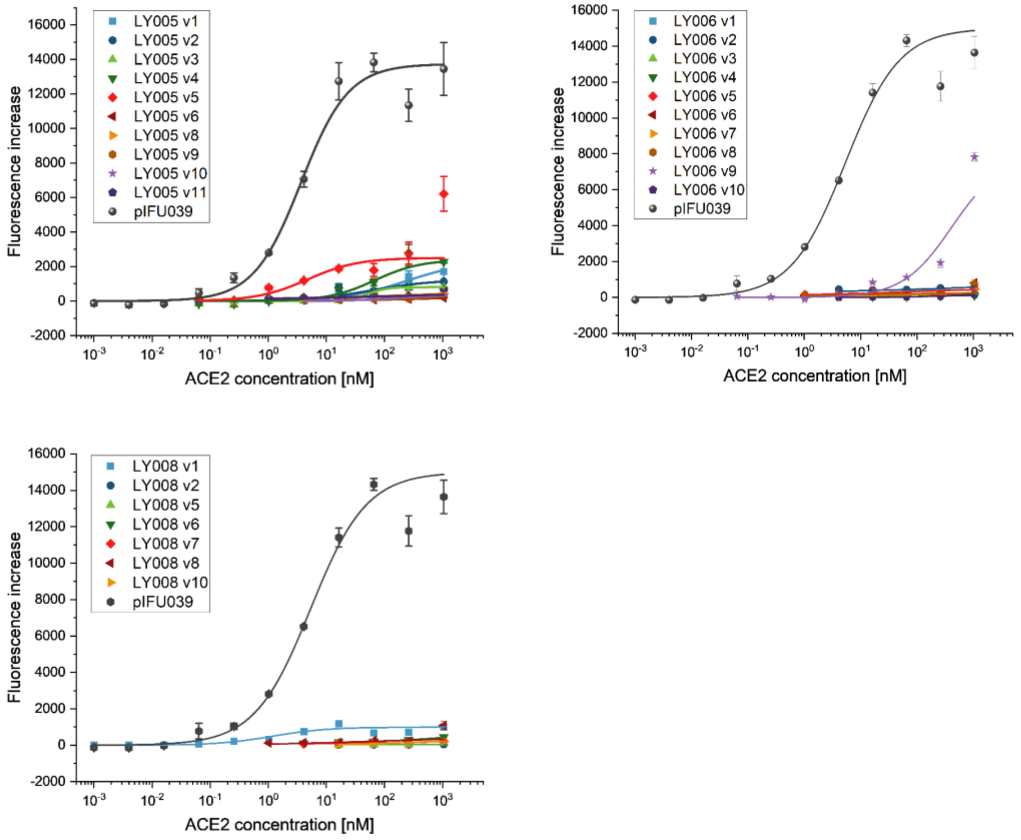


**Fig. S2.**

**Titration curves of isogenic variants.** Individual colonies were randomly selected from each library. For each variant, isogenic titrations were performed ranging from 1pM to 1μM ACE2. Titration curves are separated by library: LY005 (top left), LY006 (top right) and LY008 (bottom left). Wild type (pIFU039) titration curve is included as a control. Technical replicates were performed for each variant, error bars represent 2 s.e.m.. A wild type titration was performed in duplicate each day, therefore, error bars for pIFU039 represent 4 s.e.m.

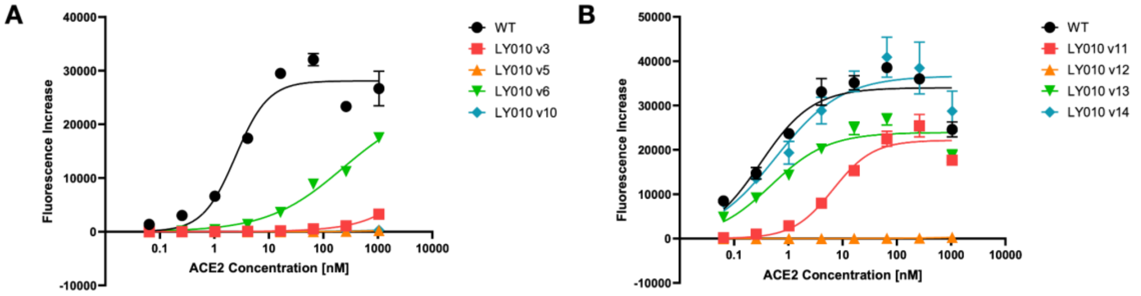

**Fig. S3. Titration curves of isogenic variants from LY010.** Individual colonies were randomly selected. For each variant, isogenic titrations were performed ranging from 1pM to 1µM ACE2. Titration curves are separated by ACE2 supplier (**A** University of Washington, Institute for Protein Design, human ACE2-Fc, Lot #20200706 **B** ab273687 ACE2, Abcam). Wild type (pIFU039) titration curve is included as a control for both ACE2 suppliers. Technical replicates were performed for each variant, error bars represent 2s.e.m.

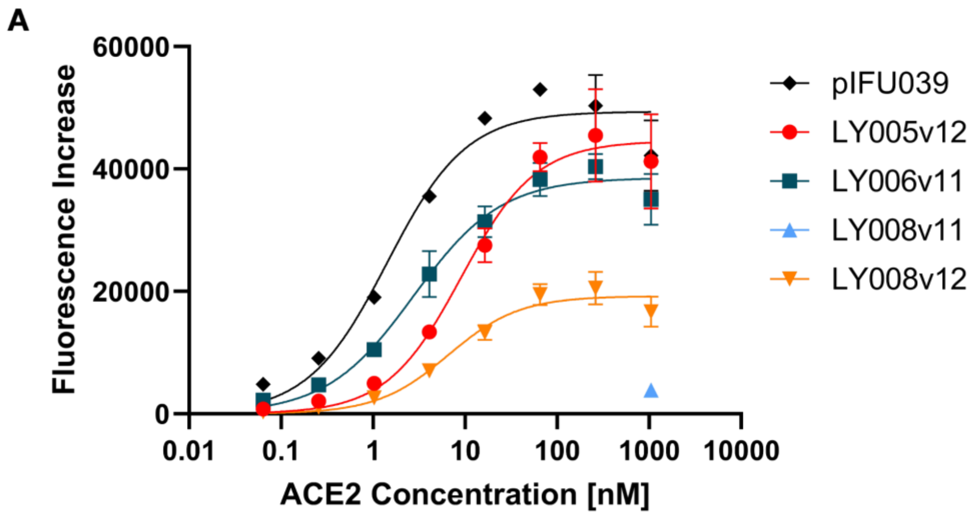


**Fig. S4. Titration curves of isogenic variants predicted to be Like-WT.** Variants were chosen to maximize mutational distance from WT and from other selected variants. For each variant except LY008v11, isogenic titrations were performed ranging from 1pM to 1µM ACE2. For LY008v11, a single binding check at 1µM ACE2 was done to demonstrate Worse-than-WT binding behavior. Wild type (pIFU039) titration curve is included as a control. Biological replicates were performed for each variant, error bars represent 2s.e.m.

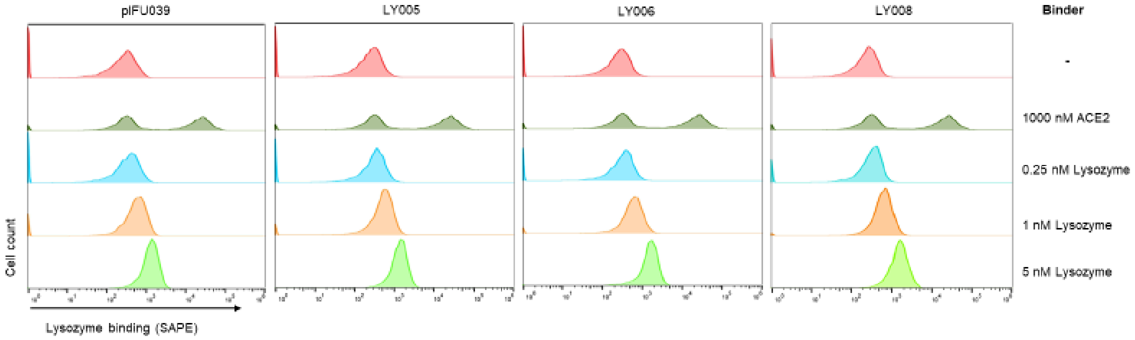


**Fig. S4 Polyspecificity assay.** The different libraries (LY005, LY006 and LY008) as well as WT (pIFU039) were labeled with different concentrations of lysozyme (0.25nM, 1nM and 5nM) for 30min at room temperature with shaking. As a control for specific binding we labeled the cells with 100nM ACE2 at the same conditions. We observed two peaks of SAPE fluorescence, one from the non-binding cells and the second from the cells displaying RBD that binds to ACE2. In the presence of lysozyme, a single peak of SAPE fluorescence is observed showing that the displayed RBD does not have non-specific binding. The fluorescence of the non-binding cells in the presence of lysozyme increases due to the binding of lysozyme to the yeast cell wall.

Variant $R_{498}P_{499}T_{500}Y_{501}Y_{505}$

$$\boldsymbol{\beta}_{1^\circ} = \boldsymbol{\beta} R_{498} + \boldsymbol{\beta} P_{499} + \boldsymbol{\beta} T_{500} + \boldsymbol{\beta} Y_{501} + \boldsymbol{\beta} Y_{505}$$

$$\boldsymbol{\beta}_{2^\circ} = \boldsymbol{\beta} R_{498}\, P_{499} + \boldsymbol{\beta} R_{498}\, T_{500} + \boldsymbol{\beta} R_{498}\, Y_{501} + \boldsymbol{\beta} R_{498}\, Y_{505} + \boldsymbol{\beta} P_{499}\, T_{500} + \ldots$$

$$\boldsymbol{\beta}_{3^\circ} = \boldsymbol{\beta} R_{498}\, P_{499}\, T_{500} + \boldsymbol{\beta} R_{498}\, P_{499}\, Y_{501} + \boldsymbol{\beta} R_{498}\, P_{499}\, Y_{505} + \boldsymbol{\beta} R_{498}\, T_{500}\, Y_{501} + \ldots$$

$$\text{Genetic Score} = \boldsymbol{\beta}_{0^\circ} + \boldsymbol{\beta}_{1^\circ} + \boldsymbol{\beta}_{2^\circ} + \boldsymbol{\beta}_{3^\circ}$$

**Figure S5:** Example of how genetic scores are computed. The variant is shown at the top. All **β**s in the expressions are unique, and refer to different states ($\boldsymbol{\beta}_{1^\circ}$), pairs of states ($\boldsymbol{\beta}_{2^\circ}$), or triplets of states ($\boldsymbol{\beta}_{3^\circ}$). The final variant score is the sum of all degrees of states in the sample, added to the global average intercept ($\boldsymbol{\beta}_{0^\circ}$).

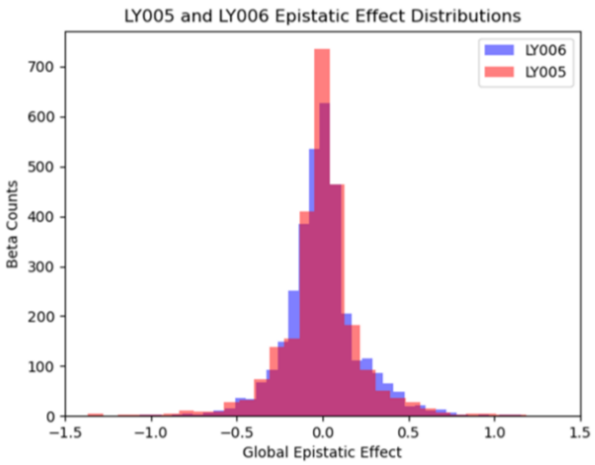


**Figure S6**: Histogram of all epistatic effect values shown in both libraries LY005 and LY006. Effects from LY005 are shown in red, and effects from LY006 are shown in blue. Main effects are excluded.

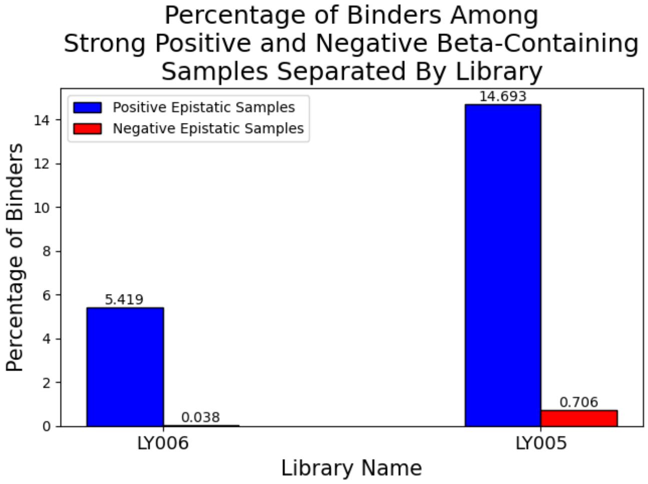


**Figure S7**: The percentage of binders observed in both LY005 and LY006 when examining samples containing strongly positive or negative epistatic terms. Samples containing epistatic effects greater than 1 are shown in blue. Samples containing epistatic effects less than -1 are shown in red.

#### Supplementary Tables

**Table S1.** Summary of Figures S2-S4.

| Sequence Name | Mutations | Titration Classification | Predicted Classification |
|---|---|---|---|
| LY005v1 | (P499L, Y501P, H505F) | worse than WT | worse than WT |
| LY005v2 | (P499F, T500W, Y501P, H505D) | NB | worse than WT |
| LY005v3 | (R498L, P499H, T500M, Y501H, H505D) | NB | NB |
| LY005v4 | (R498E, P499L, T500S, Y501T) | NB | NB |
| LY005v5 | (P499H, T500L, Y501T) | worse than WT | worse than WT |
| LY005v6 | (R498V, P499Y, H505L) | NB | NB |
| LY005v7 | (R498L, P499T, T500L, Y501P, H505F) | NB | NB |
| LY005v8 | (R498G, P499L, T500R, Y501P, H505L) | NB | NB |
| LY005v9 | (R498L, P499H, T500R, Y501P, H505L) | NB | NB |
| LY005v10 | (T500L, Y501H, H505V) | NB | NB |
| LY005v11 | (R498Q, P499F, T500L, Y501H) | NB | NB |
| LY006v1 | (F456T, Y473V, A475P, G476W, F486T) | NB | NB |
| LY006v2 | (F456T, Y473D, G476V, G485A, F486S) | NB | NB |
| LY006v3 | (F456S, Y473F, A475P, G476V, G485V) | NB | NB |
| LY006v4 | (F456S, Y473F, A475L, G476S, G485V, F486I) | NB | NB |
| LY006v5 | (F456I, Y473D, G476V, F486I) | NB | worse than WT |
| LY006v6 | (A475H, G476L, G485W, F486T) | worse than WT | NB |
| LY006v7 | (F456T, Y473L, A475P, G485L) | worse than WT | NB |
| LY006v8 | (F456I, Y473L, A475V, G476A, G485S, F486S) | worse than WT | worse than WT |
| LY006v9 | (F456I, A475V, G476S, F486T) | worse than WT | worse than WT |
| LY006v10 | (F456I, A475H, G476V, G485L) | NB | NB |
| LY008v1 | (S438A, N439K, N440D, L441K, D442Y, V445A, N450D, L452K, Y453F, R457A, K458T, N460K, K462A, F464Y, I468T, E471T) | worse than WT | worse than WT |
| LY008v2 | (S438A, N439T, N440S, L441K, D442Y, V445A, N450D, L452K, F456Y, R457A, K458T, S459R, N460K, K462A, I468T, E471T) | NB | NB |
| LY008v5 | (S438A, N439T, N440D, L441K, D442Y, V445A, N448D, N450R, L452K, Y453F, R457A, K458E, S459R, N460P, K462A, I468T, E471T) | NB | NB |

| | | | |
|---|---|---|---|
| LY008v6 | (S438A, N439T, N440D, L441K, D442Y, V445E, N450D, L452K, K458T, N460A, K462A, F464Y, R466K, I468T, E471T) | NB | NB |
| LY008v7 | (S438A, N439T, N440D, L441K, D442Y, V445A, N450D, L452K, F456Y, R457A, K458S, N460T, K462A, I468T, E471T) | NB | NB |
| LY008v8 | (S438A, N439K, N440D, L441K,D442Y, V445E, N450D, L452K, F456Y, K458T, N460A, K462A, I468T, E471T) | worse than WT | worse than WT |
| LY008v10 | (S438A, N439K, N440D, L441K, D442Y, V445A, N450D, L452K, Y453F, F456Y, R457A, K458S, N460T, K462A, I468T, E471T) | NB | NB |
| LY010v3 | (N440K, V445M, G446S, N448D, Y449R, L455Q, F456D) | worse than WT | worse than WT |
| LY010v5 | (N440K, V445N, G446S, N448T, Y449F, Y453I, F456T) | NB | NB |
| LY010v6 | (N440K, V445A, G446S, N448S, Y449F, F456A) | worse than WT | worse than WT |
| LY010v10 | (N440K, V445E, G446S, Y453E, L455V, F456P) | NB | NB |
| LY010v11 | (N440K, V445D, G446S, N448D, Y449F, Y453H, F456K) | worse than WT | worse than WT |
| LY010v12 | (N440K, V445A, G446S, N448W, Y449R, Y453D, F456T) | NB | NB |
| LY010v13 | (N440K, V445N, G446S, N448A, Y449R, Y453F, L455Q, F456T) | Like WT | Like WT |
| LY010v14 | (N440K, V445M, G446S, N448T, Y449P, Y453F, L455Q, F456I) | Like WT | Like WT |
| LY005v12 | (R498Q, P499S, T500S, Y501N, H505Y) | Like WT | Like WT |
| LY006v11 | (Y473F, A475V, G476A, G485S, F486S) | Like WT | Like WT |
| LY008v11 | (S438A, N439K, N440S, L441T, D442Y, V445A, N450D, L452K, Y453F, R457A, K458T, S459R, N460K, K462A, F464Y, I468T, E471T | worse than WT | Like WT |
| LY008v12 | (S438A, N439K, N440D, L441K, D442Y, N450D, L452K, Y453F, F456Y, K458E, S459K, N460P, K462E, F464Y, R466K, I468T, E471T) | Like WT | Like WT |

**Table S2**: Various performance metrics attained by a variety of simple machine learning architectures during the baseline assessment. BA is balanced accuracy, MCC is Matthews correlation coefficient, ACC is overall accuracy, F1 is F1 score, AUC is Area Under the ROC Curve, and AUPRC is Area Under the Precision-Recall Curve.

| Model | Library | BA | MCC | Acc | F1 | AUC | AUPRC |
|---|---|---|---|---|---|---|---|
| SVC | LY005 | 78.9 | **0.59** | **96.0** | **0.61** | **0.95** | 0.68 |
| RF | | 63.7 | 0.42 | 95.5 | 0.40 | 0.94 | 0.55 |
| KNN | | 75.5 | 0.46 | 94.0 | 0.49 | 0.89 | 0.41 |
| NB | | 56.7 | 0.36 | 95.4 | 0.24 | 0.93 | 0.68 |
| Log Reg | | **88.4** | 0.5 | 89.9 | 0.48 | **0.95** | **0.72** |
| SVC | LY006 | 50.0 | 0.00 | 99.0 | 0.00 | 0.99 | 0.95 |
| RF | | 53.9 | 0.28 | 99.1 | 0.15 | 0.78 | 0.35 |
| KNN | | 56.6 | 0.36 | 99.2 | 0.23 | 0.74 | 0.25 |
| NB | | 50.0 | 0.00 | 99.0 | 0.00 | **1.00** | **0.96** |
| Log Reg | | **63.1** | **0.49** | **99.3** | **0.41** | 0.99 | 0.67 |
| SVC | LY008 | 89.7 | **0.75** | **93.4** | **0.78** | 0.95 | 0.84 |
| RF | | 81.5 | 0.56 | 87.4 | 0.62 | 0.85 | 0.45 |
| KNN | | 73.6 | 0.46 | 86.5 | 0.54 | 0.84 | 0.42 |
| NB | | 63.9 | 0.48 | 89.3 | 0.43 | **0.99** | **0.95** |
| Log Reg | | **91.9** | 0.74 | 92.5 | **0.78** | 0.97 | 0.86 |

**Table S3:** Various performance metrics attained by logistic regression using different feature sets. BA is balanced accuracy, MCC is Matthews correlation coefficient, ACC is overall Accuracy, F1 is F1 score, AUC is Area Under the ROC Curve, and AUPRC is Area Under the Precision-Recall Curve.

| Feature Set | Evaluation Set | BA | MCC | Acc | F1 | AUC | AUPRC |
|---|---|---|---|---|---|---|---|
| One Hot | Validation (20% LY008) | 93.5 | 0.86 | 96.6 | 0.88 | **0.99** | **0.96** |
| EMME | | 52.6 | 0.12 | 85.5 | 0.12 | 0.87 | 0.43 |
| One Hot + EMME | | **95.6** | **0.89** | **97.2** | **0.91** | **0.99** | **0.96** |
| ESM2 | | 88.2 | 0.75 | 93.7 | 0.79 | 0.98 | 0.92 |
| One Hot | Test (100% LY005 + LY006) | 50.0 | 0.00 | 2.2 | 0.04 | 0.65 | 0.03 |
| EMME | | 42.5 | -0.06 | **78.5** | 0.01 | 0.28 | 0.01 |
| One Hot + EMME | | 50.0 | 0.00 | 2.2 | 0.04 | 0.46 | 0.02 |
| ESM2 | | **58.1** | **0.06** | 19.8 | **0.05** | **0.74** | **0.05** |

**Table S4:** Various performance metrics attained by Logistic Regression using different combinations of one-, two- and three-body one-hot encoded features. BA is balanced accuracy, MCC is Matthews Correlation Coefficient, ACC is overall accuracy, F1 is F1 score, AUC is Area Under the ROC Curve, and AUPRC is Area Under the Precision-Recall Curve.

| Feature Set | Library | BA | MCC | ACC | F1 | AUC | AUPRC |
|---|---|---|---|---|---|---|---|
| One Body | LY005 | 89.0 | 0.80 | 98.8 | 0.88 | 1.00 | 1.00 |
| One + Two Body | | 92.7 | 0.92 | 99.2 | 0.92 | 1.00 | 1.00 |
| One + Two + Three Body | | **93.9** | **0.93** | **99.4** | **0.94** | **1.00** | **1.00** |
| One Body | LY006 | 68.4 | 0.61 | 99.4 | 0.54 | 0.88 | 0.57 |
| One + Two Body | | 78.7 | 0.56 | 99.1 | 0.56 | **0.99** | 0.68 |
| One + Two + Three Body | | **78.9** | **0.76** | **99.6** | **0.73** | **0.99** | **0.93** |

**Table S5:** The average difference between the total score ΔΔGs of the double sample and the two single samples from **Figure 3D,** decomposed by index and energy term. The top five contributors to the double sample's enhanced energetic fitness are shown.

| Position | Score Term | Difference |
|---|---|---|
| 476 | fa_dun | -1.738 |
| 475 | fa_sol | -1.632 |
| 476 | fa_sol | -1.084 |
| 473 | fa_dun | -1.070 |
| 475 | fa_dun | -0.651 |

**Table S6:** The selected positions and allowed residues in composing libraries LY005+6. The native residue at each position is the first entry in the list of allowed residues.

| Library | Position | Allowed Residues |
|---|---|---|
| LY005 | 498 | R,Q,V,L,G,E |
| | 499 | P,F,Y,S,L,H |
| | 500 | T,W,S,R,M,L |
| | 501 | Y,T,S,N,H,P |
| | 505 | Y,F,H,V,L,D |
| LY006 | 456 | F,I,S,T |
| | 473 | Y,F,H,V,L,D |
| | 475 | A,D,V,L,H,P |
| | 476 | G,L,S,W,A,V |
| | 485 | G,A,V,L,S,W |
| | 486 | F,I,S,T |

**Table S7:** The number of variants of each binding class belonging to each library.

| Library | Total Number of Variants | Number of Like WT | Number of Worse | Number of NB | Percentage of Like WT |
|---|---|---|---|---|---|
| LY005 | 7763 | 410 | 2036 | 5317 | 5.281% |
| LY006 | 19546 | 194 | 11226 | 8128 | 0.992% |
| LY008 | 1593 | 227 | 395 | 971 | 14.250% |
| LY010 | 4587 | 1021 | 1355 | 2212 | 22.254% |

#### Supplementary Methods

##### LY008 Construction

We leveraged ProteinMPNN (1) to sample at numerous positions along the RBD to uncover sequences that are both distant from the WT and similarly functional. ProteinMPNN was used to generate sequences by sampling in cassette 2 and 3 separately. Several different design approaches were implemented. In one set, only positions which made interactions directly with ACE2 were mutated. In cassette 2, these indices were 449, 453, 455, and 456. In cassette 3, positions 475, 486, 487, 493, 496, 498, 500, 503, and 505 were targeted. In all other sets, ProteinMPNN was permitted to design the whole cassette. Additionally, a series of biologically informed biases were applied to other sequence generation runs. The Bloom lab has identified a number of point mutations strongly correlated with enhanced binding to ACE2 (2). These were E484K/Q/P, N439K, N501Y, and Y453F. Each of these 6 point mutations were individually fixed in independent ProteinMPNN runs. Additionally, designs were generated fixing all point mutations on a given cassette. For cassette 3, E484K was prioritized over E484Q and P. The GISAID database was mined to inform more designs (3). Observed point mutations across all documented viral strains were recorded. ProteinMPNN was then restricted to install either a recorded GISAID mutant residue or the native residue at the corresponding position. In combination, all of the above run restrictions and biases yielded 24 unique sequence design parameters. Finally, to ensure generated sequences would function both with and without ACE2, the native RBD was duplicated and placed 100Å away from the original complex. The two separated RBDs were then jointly sampled by tying their positions together. In this fashion, 1000 sequences were generated for each run, granting a total of 24,000 RBD designs.

Structural models were made for all designs by grafting the MPNN mutations onto the native structure (PDB ID 6M0J) using PyRosetta FastRelax. Coordinate constraints were applied to the native structure before mutation installation (4, 5). A task factory was initialized to add the relevant mutations, and the entire structure complex was relaxed. We additionally perform a round of repacking to all mutation sites and their neighbors after relaxation. The relaxation+repacking protocol is repeated five times on the same starting trajectory. The pose with the lowest total energy is accepted as the final structure for that design. The $\Delta\Delta$Gs for binding and folding were then computed. $\Delta\Delta G_{Binding}$ was determined by first separating the RBD and ACE2, repacking them, summing each chain's total score, and subtracting the summation from the total score of the complex. The native binding energy was computed in the same way and subtracted from this value. $\Delta\Delta G_{Folding}$ was calculated by taking the total score of the native RBD without ACE2, and subtracting it from the total score of the designed RBD. To supplement our calculations, we also used the established Cartesian $\Delta\Delta$G protocol to calculate additional $\Delta\Delta$Gs for binding and folding (6).

Designs were filtered down according to their computed energies, both with the PyRosetta and the Cartesian **ΔΔ**G protocols. Average **ΔΔ**Gs were computed for each metric across the entire design set. Candidate sequences were then kept in the final set if they fell below the average for all of the four measurements (PyRosetta **ΔΔ**$G_{Binding}$, PyRosetta **ΔΔ**$G_{Folding}$, Cartesian **ΔΔ**$G_{Binding}$, and Cartesian **ΔΔ**$G_{Folding}$). 1,611 designs passed this filter, which is referred to as library LY008.

**LY010 Construction**

LY010 was constructed using multiple pre-trained models to suggest functional areas in sequence space to be sampled from. To reduce the size of the search space, only particular positions were sampled for each cassette. For cassette 2, these positions were 445, 448, 449, 453, 455, and 456, all of which lie within 8Å of ACE2. Positions 444 and 457 were excluded to keep the design count at a computationally feasible level. For cassette 3, multiple position sets were sampled independently (**set 1:** positions 498-501, and 505; **set 2:** 496-507; **set 3:** 475, 476, 488, 496-507).

Three separate scoring approaches were devised to identify sequences. For the first approach, point mutations were installed individually across the whole cassette, yielding 19*N total sequences, where N is the length of the cassette. Each point mutation was scored with ESM2 relative to the Omicron variant (7, 8) and then sorted by score. The top 5 mutations at each position were retrieved, and designs were assembled by sampling them combinatorially and exhaustively. The full sequences were then scored using ProteinMPNN in scoring mode, providing the native structure in complex with ACE2 as reference (1, 2). The differences between design scores and the Omicron variant score were computed. Any samples with higher score differences than the average were discarded. To sample the sequence space efficiently, t-SNE dimensionality reduction was performed on the passing sequence set, which was further filtered based on a minimum 2-dimensional distance threshold to the rest of the set. Each of the final designs had to be at least 1 unit apart from all other accepted sequences after dimensionality reduction. For the first approach, 681 designs were generated for cassette 2, and 1,488 sequences were made for cassette 3.

For the second approach the point mutations were scored using ProteinMPNN using the native complex for reference. The Omicron variant ProteinMPNN score was subtracted from all point mutation scores (1). Mutations were sorted by score difference to select the lowest 5 at each sampled position. These mutations were then sampled combinatorially to generate full sequences, which were scored by ESM2 relative to the Omicron variant (7, 8). Sequences with negative ESM2 scores were discarded. Dimensionality reduction was conducted identically to the first approach, and designs were filtered by minimum geometric distance. The second approach generated 664 cassette 2 sequences and 1,197 cassette 3 sequences. For the third approach, the top 5 point mutations at each position as determined by both ProteinMPNN and ESM2 were combined and sampled together combinatorially. Dimensionality reduction and distance filtering was performed as previously described. This final approach generated 3,347 sequences for cassette 2 and 3,876 designs for cassette 3. Across all approaches and index sets, 4,692 new cassette 2 designs and 6,561 new cassette 3 sequences were produced. The full set of ProteinMPNN and ESM2-suggested sequences are referred to as library LY010.

**Epistasis Analysis**

Logistic regression was used for the epistatic analysis given its use in past attempts to characterize epistatic interactions (12). Feature sets were composed of either all one, two, and three-body encoded representations, only the one and two-body representations, or only the one-body terms. This partitioning allowed us to estimate the degree of binding explained by epistatic interactions between pairs and triplets of mutations. All models were trained using a 10-fold stratified split. The best performance across all folds was reported (**Figure 3A**). A full breakdown of the model performance with each feature set is shown in **Table S4**.

**Calculation of Main and Epistatic Effects**

We next identified particular samples which exhibited positive epistasis in both libraries. In line with previous efforts, we first extracted the coefficients from the full model and matched them to their corresponding mutation sets. We also retrieved the intercept of the full model, referred to here as the global average. A zero-sum constraint was then imposed on all terms, ensuring that the sum of all beta coefficients for every site and combination of sites adds up to zero. These adjusted betas were then used to compute the genetic scores of each variant corresponding to binding fitness. The genetic score of a given sequence is defined as the sum of the coefficients corresponding to all states in the sequence. In the case of library LY005, there were five mutational sites. As such, each sample would have five one-body betas associated with it. There would also be 10 two-body betas and 10 three-body betas, representing all pairs and triplets of the five positions, respectively. We also include the global average state, computed from the model intercept, and sum all terms together to obtain a variant's genetic score. An example of how genetic scores are computed is displayed in **Figure S5.** With genetic scores computed for all sequences in both libraries, we next calculated the main and epistatic effects of all coefficients of our model. The contribution of one-body betas to binding is referred to as a main effect, while the contributions of pairs and triplets are epistatic effects. A positive main or epistatic effect for a given state implies that the incorporation of said state contributes positively to binding, with more positive effects having stronger ramifications. Conversely, negative effects are expected to contribute negatively to binding activity. The main effect of a given state is defined as the average difference of the genetic scores of samples containing said state and the global average intercept. The epistatic effect of a given pair of states is similarly defined as the average difference between the genetic scores of all samples containing the target pair and the sum of the global average intercept and the constituent main effect terms of the current double-body coefficient. The epistatic effect of a triplet of states is defined in the same manner, where the sum of all constituent single and double-body betas and the global intercept is subtracted from the average genetic scores of all samples containing the target triplet.

**Energetic Validation**

To verify our epistatic model findings, we closely examined the samples that logistic regression identified as exhibiting pairwise positive epistasis. We selected two-body beta design (Y473F,A475V,G476A,G485S,F486S) from LY006, as this sequence had the largest difference in binding probability between itself and its constituent one-body beta samples. These one-body designs were (Y473F,A475V,G476W,G485S,F486S) and (Y473F,A475H,G476A,G485S,F486S). The samples (Y473F,A475V,G476A,G485S,F486S), (Y473F,A475V,G476W,G485S,F486S), and (Y473F,A475H,G476A,G485S,F486S) will be referred to as the double sample, single sample 1, and single sample 2 respectively. The pairwise epistatic effect exhibited by the double sample is (A475V,G476A). The double sample had an experimentally derived binding probability of 100%, while both single samples had probabilities of 0%, making this design set an ideal candidate for energetic validation.

We next examined the PyRosetta-generated structural models for the double sample and both single samples to determine if energetic measurements could corroborate our epistatic analysis (4). To this end, we computed the total energy of each sample across the whole RBD. These scores were subtracted from the native to obtain ΔΔG values for the single and double samples. Across all selections, the double sample consistently achieved superior ΔΔGs than both single samples, as seen in **Table S5**.

The energetics at each sampled position in LY006 were next examined to determine the precise cause of the total score discrepancy. Position selectors were initialized separately for positions 456, 473, 475, 476, 485, and 486. The total score was decomposed into its constituent energy terms, so each component could be investigated individually. Each position on the single and double samples had their respective score terms calculated. The WT values at the same positions were then subtracted from the double and single sample energies, yielding ΔΔGs for every score term in the total energy function. The double sample and single sample ΔΔGs were then subtracted to determine the score terms and positions which most contributed to the double sample's increased energetic fitness. The average of the differences between the double sample and the two single samples for each score term was calculated, and used to sort the energetic terms at each position and find the biggest contributors. The 5 largest discrepancies where the double sample is significantly more favorable than the single samples, as indicated by a lower energetic term difference, are shown in **Table S6**. The total_score score term was excluded for this analysis.

**Sequence Attention Model Architecture**

We employed an attention-based model to process sequences encoded by ESM2. The model first took as input the 2560-dimensional vector for each sequence. Layer normalization was performed, followed by self-attention encoding using Pytorch's MultiHeadAttn module (13). The output representation was again normalized before being processed by an MLP consisting of three fully connected layers which returned the final binding prediction. Tunable hyperparameters included attention dropout rate, batch size, learning rate, and the number of attention heads. All models were trained for 200 epochs with checkpoints every 25 epochs.

**Graph Attention Model Architecture**

RBD graph inputs to the model were first normalized by batch before being fed into the GATv2Conv graph operator implemented through PyTorch Geometric (14, 15). The output was then normalized again before being passed through a LeakyReLu activation function. This process was repeated across two more encoding blocks, each consisting of a graph attention layer, followed by normalization and activation. The final encoded representation was then provided to a fully connected layer for binding prediction readout **(Figure 5B)**. Tunable hyperparameters included dropout rate, batch size, learning rate, and the number of graph attention heads. All models were trained for 200 epochs with checkpoints every 25 epochs.
